# Spatiotemporal Mapping of Glomerular Endothelial Regeneration Identifies *Apold1* as an Intrinsic Regulator of Repair

**DOI:** 10.64898/2026.10.08.757579

**Authors:** Meike Seitz, Hannah Weißbach, Lilly Dubau, Christoph Daniel, Pilar Hartmann, Anika Wirth, Maria Schuster, Shila Azizolli, Akua Annoth, Irina Simonova, Anne Steglich, Thomas Conrad, Janine Altmüller, Vladimir Todorov, Christian Hugo, Jan Sradnick

## Abstract

Endothelial cell (EC) injury is central to the pathogenesis of various renal disorders, highlighting the critical importance of vascular repair for restoring organ health. However, due to limited access to human tissue during active regeneration, the exact reparative processes remain largely unknown. This study aimed to investigate regeneration mechanisms specifically within the glomerular microvasculature by utilizing a selective endothelial cell injury (ECI) model in mice. Lineage tracing identified the origin of regenerating ECs. To uncover the transcriptional dynamics driving these reparative changes, we combined single-cell RNA sequencing and Xenium spatial transcriptomics (ST), mapping distinct glomerular EC (gEC) subpopulations across different phases of injury and recovery. Apold1 emerged as a key regulatory candidate within this process. To functionally validate this gene’s specific role during the regenerative phase, Apold1-deficient (Apold1-/-) mice underwent the ECI model. After showing that gECs regenerate primarily from local, pre-existing renal ECs, our transcriptomic analysis identified 10 distinct gEC subpopulations and revealed a complex temporal pattern of regeneration. This process begins with the expression of genes associated with inflammation, followed sequentially by endothelial cell (EC) migration, the activation of immediate early genes, vascular remodeling and cellular proliferation. Within this dynamic process, the EC-specific gene *Apold1* was identified as a key regulatory candidate. *Apold1^-/-^* mice exhibited impaired endothelial regeneration, characterized by increased endothelial activation and greater immune cell infiltration seven days post-ECI. To our knowledge, this is the first study to elucidate the spatiotemporal mechanisms of glomerular endothelial cell regeneration. By identifying local gECs as the primary source of repair, and stablishing the change in their expression pattern during regeneration, as well as identifying *Apold1* as an intrinsic regulator, we have laid the groundwork for future research on renal microvascular disorders.

**Translational Statement:** Using EC-lineage-tracing mice and transcriptomic-techniques, we identified new endothelial subpopulations that drive injury and regeneration post-ECI. We provide insights into the local processes that orchestrate gEC regeneration and highlight complex gene patterns with several newly identified targets in renal endothelial regeneration (e.g. immediate early genes and *Apold1*). These results hold strong potential for clinical translation stimulating the development of cell therapies or other strategies to support the physiological endothelial regeneration. We functionally validate the gene *Apold1* in the process of gEC regeneration. Future steps include thoroughly characterizing the influence of *Apold1* during endothelial regeneration in mice and humans, and exploring its clinical application to improve endothelial recovery in microvascular renal diseases.

## INTRODUCTION

In the kidney, endothelial cells (ECs) play an essential role in regulating blood flow, glomerular filtration, tubular function, and the delivery of nutrients and oxygen to all renal cell types^1^. This is particularly relevant in glomerular capillaries, where fenestrated glomerular endothelial cells are in direct contact with the blood and are essential for filtration. Given these central functions, EC injury (ECI) is a key feature of almost all renal disorders (e.g. diabetic and hypertensive nephropathy, vasculitis, ischemia/reperfusion injury or IgA nephropathy^2,3^). In thrombotic microangiopathies (TMAs) or typical/atypical hemolytic uremic syndrome (HUS), severe primary ECI can lead to acute or chronic life-threatening organ damage, with the kidney being particularly affected. The introduction of therapies, such as complement inhibition in atypical HUS patients, has demonstrated that renal EC regeneration in human disease can occur^4^.

However, the mechanisms driving EC repair remain poorly understood, largely due to limited access to human tissue during regeneration phases given the risks associated with repeated kidney biopsies. Thus, experimental animal models are crucial for examining these processes. We have developed one of very few animal models mimicking human TMA^5^, enabling reproducible monitoring of glomerular ECI and regeneration within a well-defined time window to closely investigate glomerular EC (gEC) injury and repair mechanisms. For decades, bone-marrow-derived endothelial progenitor cells have been proposed to contribute to endothelial repair^6^, yet direct evidence remains limited^7^. In contrast, our prior experiments demonstrated that endothelial regeneration in murine kidneys depends exclusively on local renal mechanism^8^, and excluded renin lineage cells as a source of regenerating ECs^9^. Thus, we and others suspect local ECs to be highly involved in the process of regeneration^10,11^. Furthermore, while previous single-cell RNA sequencing (scRNAseq) studies have revealed substantial phenotypic heterogeneity among renal ECs and identified diverse subtypes within the kidney^12^, to our knowledge no scRNAseq studies have focused specifically on the molecular mechanisms driving renal gEC regeneration.

Building on these observations, we used scRNAseq and Xenium spatial transcriptomics to identify and characterize distinct endothelial subpopulations that emerge dynamically during tissue injury and subsequent regeneration in our animal model. This high-resolution approach identified candidate regulators of endothelial repair and highlighted the endothelial-specific gene *Apold1*, a known regulator of pathological angiogenesis^13^, whose role in the kidney has remained undefined. To assess the functional implications of *Apold1* in renal gEC regeneration, we conducted an evaluation of the response of *Apold1*-deficient mice (*Apold1*^−/−^) to selective gEC injury.

## METHODS

### Experimental animals

All animal experiments were conducted at the *animal facility* of the medical faculty at TUD Dresden University of Technology and complied with German animal welfare laws and were approved by local authorities (DD24-5131/394/34; DD25-5131/680/18). Male mice, aged 8- to 12-weeks, were group-housed and had access to water and food *ad libitum*. Upon sacrifice, all mice were anesthetized using ketamine/xylazine (10mg/ml ketamine, 1 mg/ml xylazine, 10 ml/kg body weight), followed by cardiac perfusion.

*Cdh5(PAC)-ERT2-tdTomato* reporter mice (provided by the Boor Lab, RWTH Aachen) for lineage-tracing experiments were induced by tamoxifen administration (concentration: 20mg/ml; 60 µl/10 g body weight) for five days, followed by a washout period. Partial biopsies were taken from the right kidney to assess recombination efficiency 24 hours before site-selective endothelial injury was induced in the left kidney as described previously^8,9^ (Supplementary Figure S1a). Final biopsies were taken 7 days after injury). Renal tissue was processed for flow cytometry as described previously^8^.

*Tg(TIE2GFP)287Sato/J* (Charles River, Germany), and *C57BL/6 J-Apold1tm1(LacZ)* mice (*Apold1^-/-^*, provided by Johannes Bohacek/ Katrien de Bock, ETH Zürich) are mouse strains with constitutive alterations.. *C57BL/6JRj* mice are used for the ST approach and as WT controls for *Apold1^-/-^.* ECI was induced in all mouse strains in the left kidney, and final biopsies of the left kidney were taken 1, 2, 4, 7 and/or 14 days post-injury (Supplementary Figure S2a). If there had been complications during the surgery procedure, mice were sacrificed to reduce animal suffering and the tissue was excluded from the analysis. An overview about the setups and animal numbers for all experiments can be found in the supplement (Supplementary Figure S2a).

### Renal endothelial cell isolation and sorting

EC isolation for scRNAseq was done in *Tg(TIE2GFP)287Sato/J* mice according to the protocol of Dumas and colleagues^14^. Briefly, renal tissue was minced, digested, and then filtered sequentially through a 40 µm filter to isolate glomeruli (>80% purity). The collected glomeruli were further dissociated. The single-cell suspension was then blocked with an Fc-quenching antibody and cells were stained with antibodies directed against CD102, CD105, CD73 and CD45 (Supplementary Table S1). The separation was done via fluorescence-activated cell sorting (FACS) and only cells with the following phenotype were further used for scRNAseq: Tie2-EGFP+, CD102+, CD105+, CD73-, CD45-.

### Single-cell RNA sequencing

Single-cell suspensions of FACS-sorted gECs were processed using the Chromium Single Cell 3′ Library platform (10x Genomics Chromium Single Cell platform) to generate barcoded scRNA-seq libraries (Supplementary Figure S2a). The resulting libraries were sequenced on an Illumina platform using paired-end sequencing. Raw data were preprocessed and demultiplexed using Cell Ranger (version 6.1.2). Quality control and analysis was performed using Seurat^15^ (version 5) for R (version 4.2.2) with RStudio (version 2026.08.2)^16^. Cells were filtered based on the following criteria: cells < 100 or > 3000 gene counts or cells with > 10 % mitochondrial counts were removed. Cells were checked for EC marker *Cdh5* and gEC marker *Ehd3* (Supplementary Figure S3a). Contaminating cells expressing markers for mesangial cells (*Des*, *Itga8*), podocytes (*Wt1*, *Nphs2*), or erythrocytes (*Hbb-bt*, *Hba-a2*) (Supplementary Figure S3b) were also removed, leaving 23 586 high-quality gECs. A total of 32 524 genes were detected. Data were normalized using the *NormalizeData*^15^ function of Seurat. Variable genes were identified using the *FindVariableFeatures*^15^ function which returned 2000 variable genes per dataset. Next, a linear transformation is applied by the *ScaleData*^15^ function with the default parameters to prevent the domination of highly expressed genes in downstream analyses.

### Spatial transcriptomics

Formalin-fixed, paraffin-embedded kidney tissue was collected and cut into 4 µm thick sections. Sections were run on the 10x Genomics Xenium platform with the predesigned mouse multi-tissue v1 (mMulti-tissue) panel and a custom panel consisting of the top genes identified in our scRNAseq analysis (Supplementary Figure S2b). Raw data were preprocessed and demultiplexed using *Space Ranger*. Quality control and analysis were performed with Seurat (version 5) and the Xenium Explorer (version 3.0.0) with data from all tissues merged for gene expression analyses. Normalization was conducted using *sctransform*^17^ (version 0.4.1).

### Bioinformatic data analyses

#### Graph-based cluster analysis

Graph-based cluster analysis was carried out on scRNAseq data. A principal component analysis (PCA) was applied to the variable genes identified earlier. Using input of the PCA, a KNN graph was constructed with the *FindNeighbors*^15^ function in Seurat. Cells were clustered according to their gene expression profiles using *FindClusters*^15^ function via the Louvain algorithm and visualized in a Uniform Manifold Approximation and Projection (UMAP) plot. Clusters were characterized using the *FindAllMarkers*^15^ function of Seurat to list all cluster-defining differentially expressed genes (DEGs) and rank them according to the log2 fold change. The top 50 genes from each cluster were reviewed through literature research for classification of the cluster (Supplementary Table S2). DEGs of our scRNAseq data, was visualized with Seurat’s *DoHeatmap*^15^ function. Genes were ranked by their log2 fold change and plotted by cluster and time points. For clarity, gene expression was averaged per time point using the *AverageExpression*^15^ function. Compositional analyses were calculated in R using the packages *dplyr* ^18^ and visualized in a stacked barplot.

#### Gene Set Enrichment Analysis

Gene set enrichment analyses (GSEA) was performed using the *clusterProfiler*^18^ package (version 4.2.2) for R. The top 100 genes from the cluster of interest was compared with the Gene Ontology biological process gene set (M5, GO:BP) from the Mouse Molecular Signatures Database (MSigDB). The results (Supplementary Table S3) were summarized by removing redundant GO terms with the open-source website revigo ^19^ and visualized as bar plots.

#### Gene expression analysis

Gene expression levels in scRNAseq and ST data were calculated using *rowMeans* ^15^ function of Seurat separately for each replicate per time point and visualized for each gene of interest as the normalized expression over time.

#### Analysis of Xenium spatial transcriptomics data

Glomeruli on the kidney sections were analyzed using Xenium Explorer (version 3.0.0) and Seurat Glomerular area was identified by the expression of markers *Itga8*, *Edh3* and *Wt1* and was manually selected to obtain glomerular cell IDs. Respective cell IDs were extracted from the whole kidney section data using Seurat. The glomerular area was then filtered for cells expressing 2 to 4 of the top DEG of the scRNAseq clusters in combination with the EC marker *Kdr* by using the *subset* function of Seurat. The used markers are specified in Supplementary Table S4. Identified cells were mapped again in Xenium Explorer and quantified by counting the number of cells per glomerulus.

### Histology

Mouse kidney tissues were fixed in zinc fixative solution (0.5 g calcium acetate, 5 g zinc acetate, 5 g ZnCl2 dissolved in 0.1 M TRIS-buffered solution (pH 7.4) for 24 hours at 4°C, embedded in paraffin, and cut into 2 µm sections. Periodic acid–Schiff (PAS) staining was performed to evaluate glomerular microthrombi ^5,20^. Fluorescence staining was conducted with antibodies against EC marker ERG, anti-RFP (tdTomato), and leukocyte marker CD45 as described previously. IHC staining in mouse tissue was done for EC marker EMCN as previously described ^20^ (Supplementary Table S1). For evaluation, the marker-positive area was assessed using computer-assisted image analysis as described earlier^21^. IHC staining of human formalin-fixed and paraffin-embedded kidney tissue was performed using antibodies against Atf3, Egr1, Serpina3, Dnajb1, Bst2, Sox17. Additional methodological details are provided in the Supplementary Methods.

### Quantification and Statistical Analysis

Analysis was done in a blinded-manner with the researcher being unaware of the group allocations. Statistical analyses were conducted using GraphPad Prism (version 10.2.3), with data presented as scatter dot plot with means ± SD. Comparisons between two groups were done via Welch’s t-test. Comparisons across multiple groups were analyzed using one-way ANOVA followed by Tukey’s post-hoc test. A p-value < 0.05 was considered statistically significant.

## RESULTS

### Single-cell transcriptomic analyses of gEC reveal clear patterns of injury followed by regeneration

We first investigated whether a distinct extra-endothelial niche contributes to endothelial cell regeneration after ECI. Following ECI, there was a marked loss of glomerular endothelial cells on day 1, followed by substantial repair by day 7 (Figure 1a-c)), and lineage tracing in inducible Cdh5(PAC)-ERT2 tdTomato reporter mice demonstrated that regenerated renal ECs arise from pre-existing adult ECs (Supplementary Figure S 1b-d). Using inducible Cdh5(PAC)-ERT2 tdTomato reporter mice, we showed that the proportion of tdT+/ERG+ cells remained consistent across all time points, indicating that regenerated ECs arise exclusively from labeled ECs persisting after injury. Thus, the regeneration of renal ECs predominantly originates from pre-existing adult renal tdTomato-positive ECs. In order to further investigate the mechanisms in the local ECs during injury and regeneration, gECs undergoing the ECI model were isolated and characterized using fluorescence-activated cell sorting and single-cell transcriptomics (Figure 2a, Supplementary Figure S2b).

**Figure 1.**
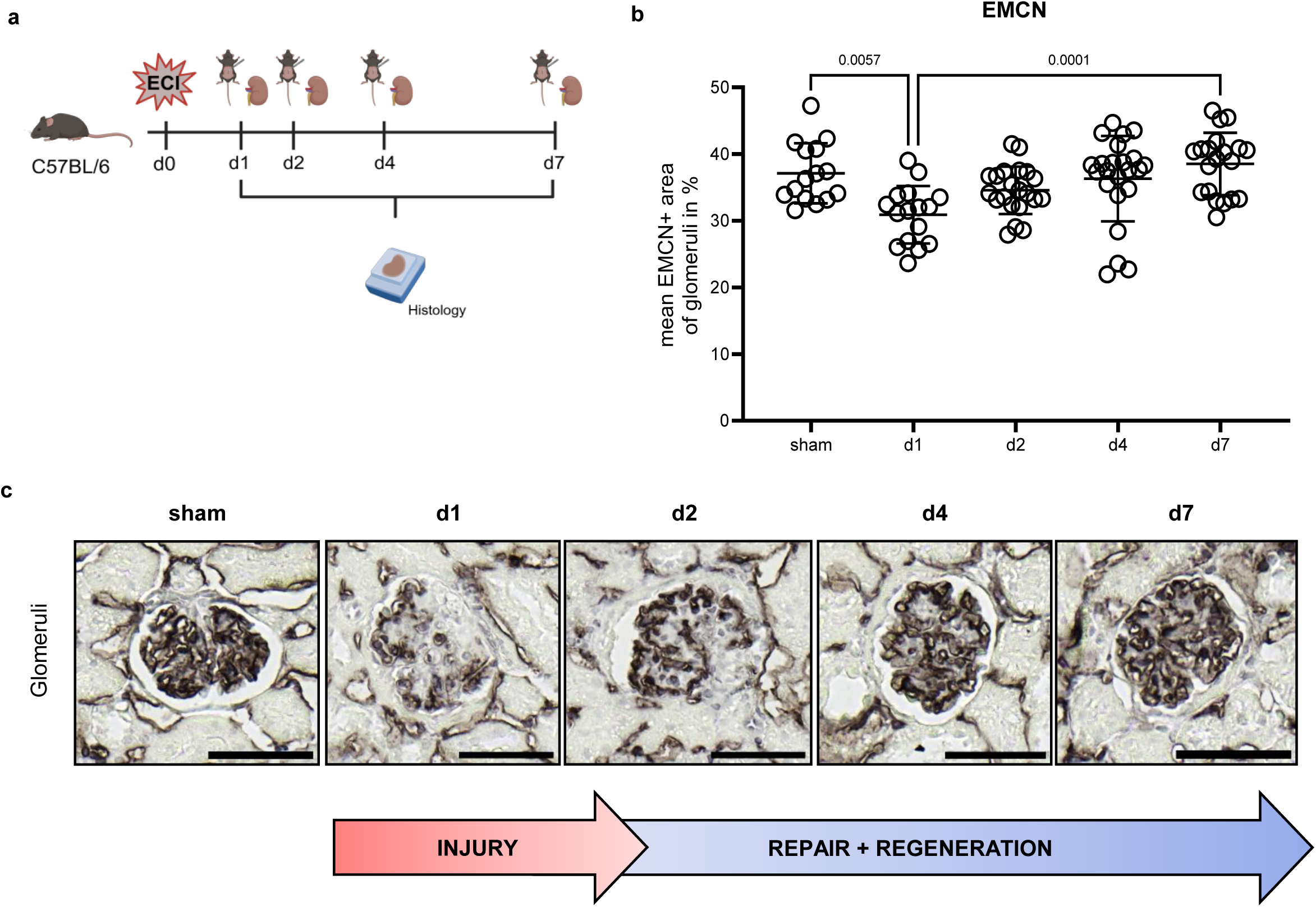
Renal endothelial cells are predominantly regenerated by an adult renal endothelial cell pool. (a) Experimental setup to follow injury and regeneration of endothelial cells after the endothelial cell injury model in *C57BL7/6* mice. Kidney biopsies for histology are taken 1 (d1), 2 (d2), 4 (d4), and 7 (d7) days after injury induction (d0). Image created in BioRender. (b) Percentage of mean EMCN*-*positive glomerular area in IHCkidney tissue of sham and ECI-mice. Data are shown as scatter dot plot with means ± SD. (c) Representative images of EMCN-positive area in mouse kidney tissue, representing the course of the ECI model in the glomerular area. Scale bar: 50µm

**Figure 2.**
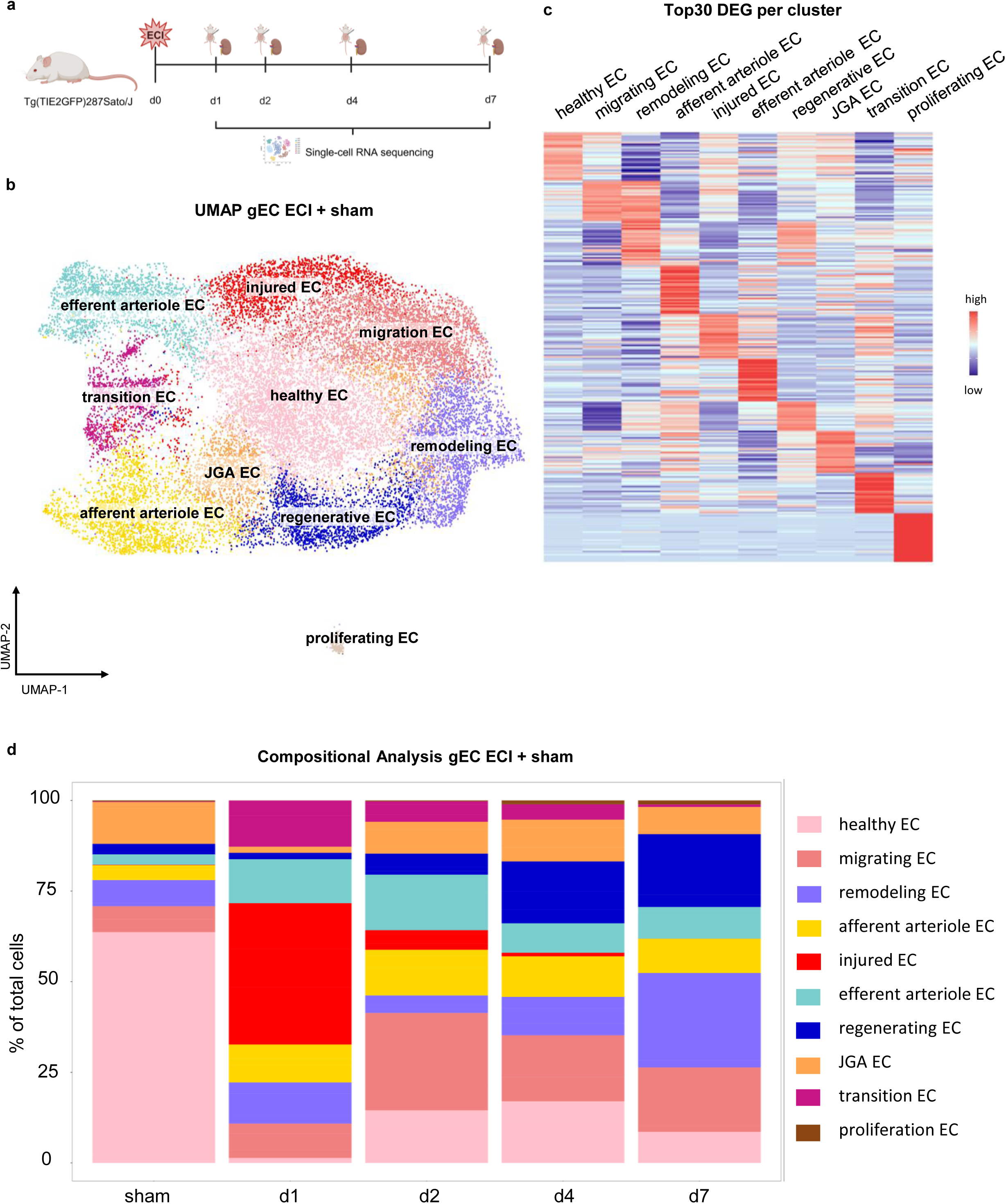
Single-cell transcriptomic analyses of glomerular endothelial cells reveal clear patterns of injury followed by regeneration. (a) Experimental setup for the single-cell RNA sequencing approach. *Tg(TIE2GFP)287Sato/J* mice are induced with the endothelial cell injury model (d0). Kidney biopsies are taken on days 1 (d1), 2 (d2), 4 (d4), and 7 (d7) after injury, and isolated glomerular endothelial cells are analyzed with 10x Chromium after further preparation as described inSupplemental Figure 1b. (b) UMAP visualization of glomerular endothelial cell cluster analysis from sham and ECI mice revealed 10 different endothelial cell clusters. (c) The scaled heatmap with averaged expression of single-cell RNA sequencing data of the top 25 differentially expressed genes for each cluster of the glomerular endothelial cells. (d) The compositional analysis shows the proportions of each cluster of the total number of cells for each time point. Colors for each cluster are equal to the UMAP plot. DEG = differentially expressed genes; gEC = glomerular endothelial cells; ECI = endothelial cell injury; UMAP = Uniform Manifold Approximation and Projection

Cluster analysis of sham-operated gECs unveiled seven clusters corresponding to markers of gECs of afferent/efferent arteriole, glomerular capillaries, and proliferating ECs (Supplementary Figure S4a). During the ECI and regeneration phases, the number of EC clusters varied per time point (Supplementary Figure S4b). To investigate whether clusters recur or are novel throughout the experiment, we performed a Jaccard similarity analysis of each single time point compared to each other (Supplementary Figure S4c). This analysis revealed the appearance of unique gEC clusters, potentially involved in the injury and regeneration processes observed throughout our ECI model.

The graph-based analysis of all gECs together identified ten distinct clusters, which were visualized in a UMAP plot (Figure 2b). Using Seurat’s *FindAllMarkers* function, we identified the top 30 cluster-defining DEGs demonstrating high uniqueness for each group (Figure 2c, Supplementary Table S2). The cell populations were classified using literature research based on the top 50 DEGs per cluster (Supplementary Table S2).

The largest cluster comprises *healthy EC*s expressing capillary markers related to fundamental physiological processes^12,22–24^, representing the highest proportion of cells in the healthy sham group. This proportion decreases on day 1 post-ECI before expanding again toward day 7, as expected with our model (Figure 2d). Cluster representing ECs from morphological structures such as *afferent*^12,25^/*efferent*^12,26^ *arteriole* and *JGA*^12,27^ *ECs* show only slight temporal variations (Figure 2d).

On day 1 post-ECI, an *injured EC* subset emerges, defined by genes regarding inflammation and immune system activation^28–31^. Nearly absent in sham mice, this group peaks significantly on day 1 and returns to baseline levels by day 7 (Figure 2d). The *transition EC* phenotype exhibits similar kinetics, peaking on day 1 and decreasing through day 7. Characterized by regenerative^32–34^ (Supplementary Table 3) and injury-related genes^35,36^, these cells potentially represent an intermediate stage towards the repair phase. The cluster of *migrating ECs* peaks on day 2 post-ECI, expressing genes related to cell adhesion and reorientation^37–42^ (Figure 2d). Subsets associated with later regenerative stages, such as *remodeling* and *regenerative ECs*, increase substantially on days 4 and 7 (Figure 2d). The *remodeling EC* group is characterized by activation or cell stress markers with subsequent remodeling^43–48^ properties, while the cluster *regenerative EC* features high numbers of immediate early genes (IEGs) and genes linked to regenerative processes ^49–53^ ^51,54^. Finally, the smallest cluster, *proliferating ECs*^55–58^, is minimal in sham mice but demonstrates a robust increase during the later phase of the process on days 4 and 7 (Figure 2d).

### The progression and restoration of endothelial cell injury can be spatially verified

To examine the spatial distribution of gEC clusters during injury and regeneration, we utilized high-resolution ST on the Xenium platform by 10x Genomics (Supplementary Figure 2b; Figure 3a). Transcript density maps of the injury markers *C3* and *Bst2*, and regeneration-associated marker *Egr1*, provided an overview of injury and regeneration within our ECI model (Supplementary Figure S5a). *C3* and *Bst2* showed increased expression on day 1 post-injury, returning to baseline by day 7, while *Egr1* expression peaked on days 4 and 7, which aligns with the expected timeframe for regenerative processes seen in the scRNAseq data (Supplementary Figure S5b). The scRNAseq and ST data demonstrated a high concordance in gene expression patterns across the glomerular area, confirming consistency between these methods (Supplementary Figure S5c).

**Figure 3.**
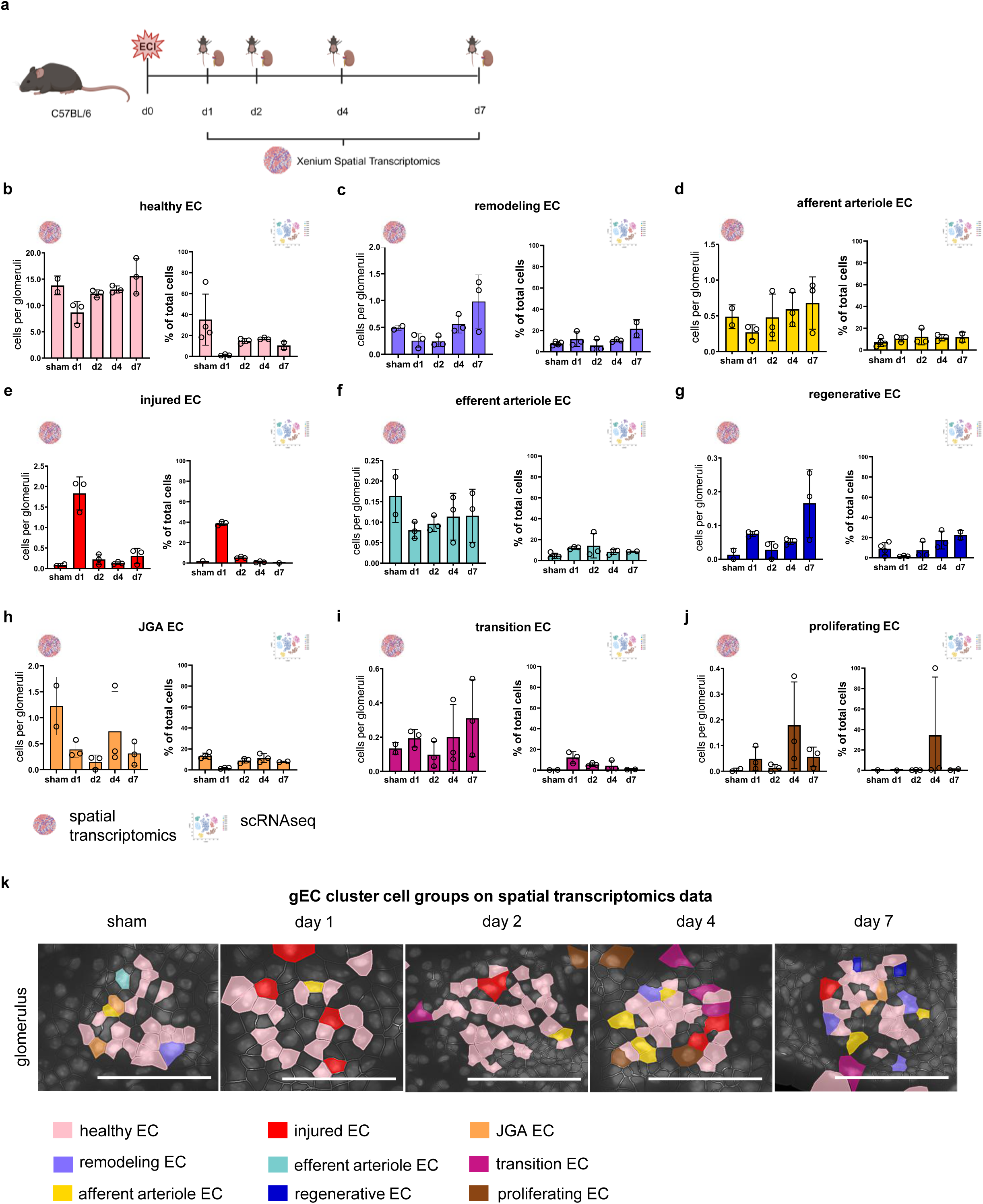
The progression and restoration of endothelial cell injury can be spatially verified. (a) Experimental setup for the spatial transcriptomics approach. *C57BL7/6* mice are induced with the endothelial cell injury model (d0). Kidney biopsies are taken on days 1 (d1), 2 (d2), 4 (d4), and 7 (d7) after injury, and PFA-fixed, paraffin-embedded tissue sections are run with 10x Xenium (as described in Supplemental Figure 1c). Image created in BioRender. (b-j) To the left of each index is the quantification of the number of cells associated with the respective cluster per glomerulus in spatial transcriptomics. On the right is the proportion of cluster-associated cells at each time point in single-cell RNA sequencing. Data are presented as mean ± SD. (t) Cutout images of renal sections with cell segmentation from 10x Xenium platform for every timepoint in glomerular area. Colored cells represent cells with the presence of the respective EC-cluster and the EC marker *(Plat and Cdh5)*., scale bar = 50 µm, gEC = glomerular endothelial cell; JGA = juxtaglomerular apparatus; EC = endothelial cells; scRNAseq = single-cell RNA sequencing

To spatially map the gEC subtypes within the glomerular area over time, we designed a custom panel based on the top scRNAseq DEGs for each cluster. We then filtered the ST data within the glomerular area to identify cells co-expressing the general EC marker *Kdr* alongside two to four of the top DEGs defining each scRNAseq cluster (Supplementary Table S4). The cluster of *migrating EC* was excluded from this analysis due to a lack of robust markers in the panel.

Quantification of cells per gEC cluster in the ST data revealed temporal changes that closely mirrored those seen in the scRNAseq data, demonstrating a comparable development of injury and regeneration processes (Figure 3b-j). Spatial mapping revealed that afferent/efferent arterioles and JGA-associated ECs mostly localized to their expected areas, while inflammatory and regenerative clusters displayed a heterogeneous glomerular distribution (Figure 3k).

### Selected scRNA-seq-derived markers show altered glomerular expression in human kidney biopsies from patients with thrombotic microangiopathy

To assess the relevance of the identified transcriptomic markers in human thrombotic microangiopathy (TMA)-associated kidney disease, we validated selected candidate proteins involved in injury, regeneration, and remodeling by immunohistochemistry. The selected markers included SERPINA3, ATF3, DNAJB1, EGR1, BST1, and SOX17, all of which showed glomerular staining in kidney biopsies from patients with, complement-mediated TMA, malignant hypertension-associated TMA and minimal change disease (MCGN), that served as controls (Supplementary Figure S6).

Patients with complement-mediated TMA and malignant hypertension-associated TMA had significantly higher serum creatinine levels, inflammatory index scores, and segmental sclerosis than patients with minimal change diseases (MCGN) (Supplementary Table S6). Therefore, minimal change disease samples were used as control group.. Quantification showed thatmalignant hypertension-associated TMA, compared to controls, had significantly increased glomerular staining for SERPINA3 and SOX17, (Figure 4a;b) as well as a higher number of EGR1-positive cells (Figure 4b). The ATF3-positive glomerular area was also significantly increased in this group. In complement-mediated TMA, the DNAJB1-positive glomerular area was significantly elevated (Figure 4c). By contrast, BST2, a marker associated with endothelial cell remodeling, did not differ significantly between the TMA groups and controls (Figure 4a).

**Figure 4.**
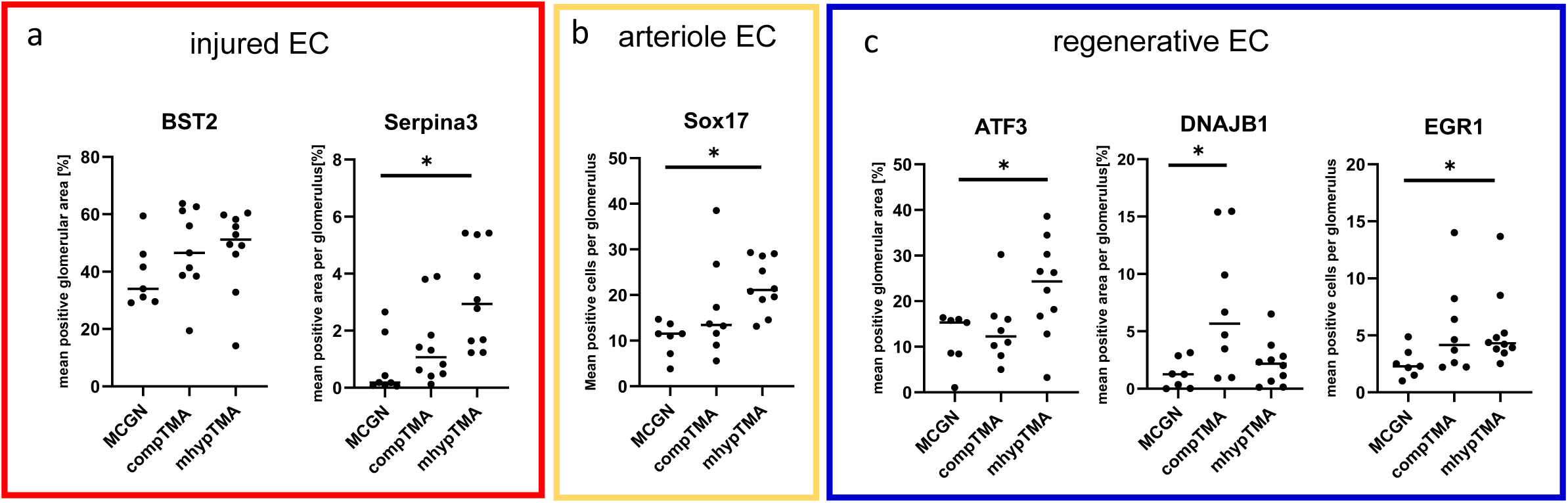
Immunohistochemical evaluation of kidney biopsies from patients with minimal change glomerulonephritis, complement-mediated thrombiotic microangiopathy, and malignant hypertension-associated thrombiotic microangiopathy,. Quantification of the mean glomerular area positive after immunohistochemical staining with antibodies against (a) BST2 and SERPINA3, (b) SOX17, and (c) ATF3, DNAJB1, and EGR1 in kidney biopsies from patients with MCGN, complement-mediated TMA, or malignant hypertension-associated TMA. Statistical comparisons were performed pairwise between the MCGN group and each TMA group. MCGN, minimal change glomerulonephritis; TMA, thrombotic microangiopathy.

### Apold1 is associated with regenerative endothelial states in mice and increased expression in human kidney disease

Both scRNAseq and ST analyses showed several endothelial subpopulations playing a potential role in the regeneration phase. To further evaluate the biological relevance of the detected genes underlying these clusters, we first focused on the *regenerative EC* cluster. The GSEA, which assesses collective shifts in pathway activity of cells, revealed the MAPK cascade, the ERK1/ERK2 pathway, and angiogenetic processes as highly enriched within this cluster (Figure 5a). A recurring gene present not only in several top-enriched gene sets (Figure 5a, red boxes) but across multiple other pathways was the gene *Apold1*.

**Figure 5.**
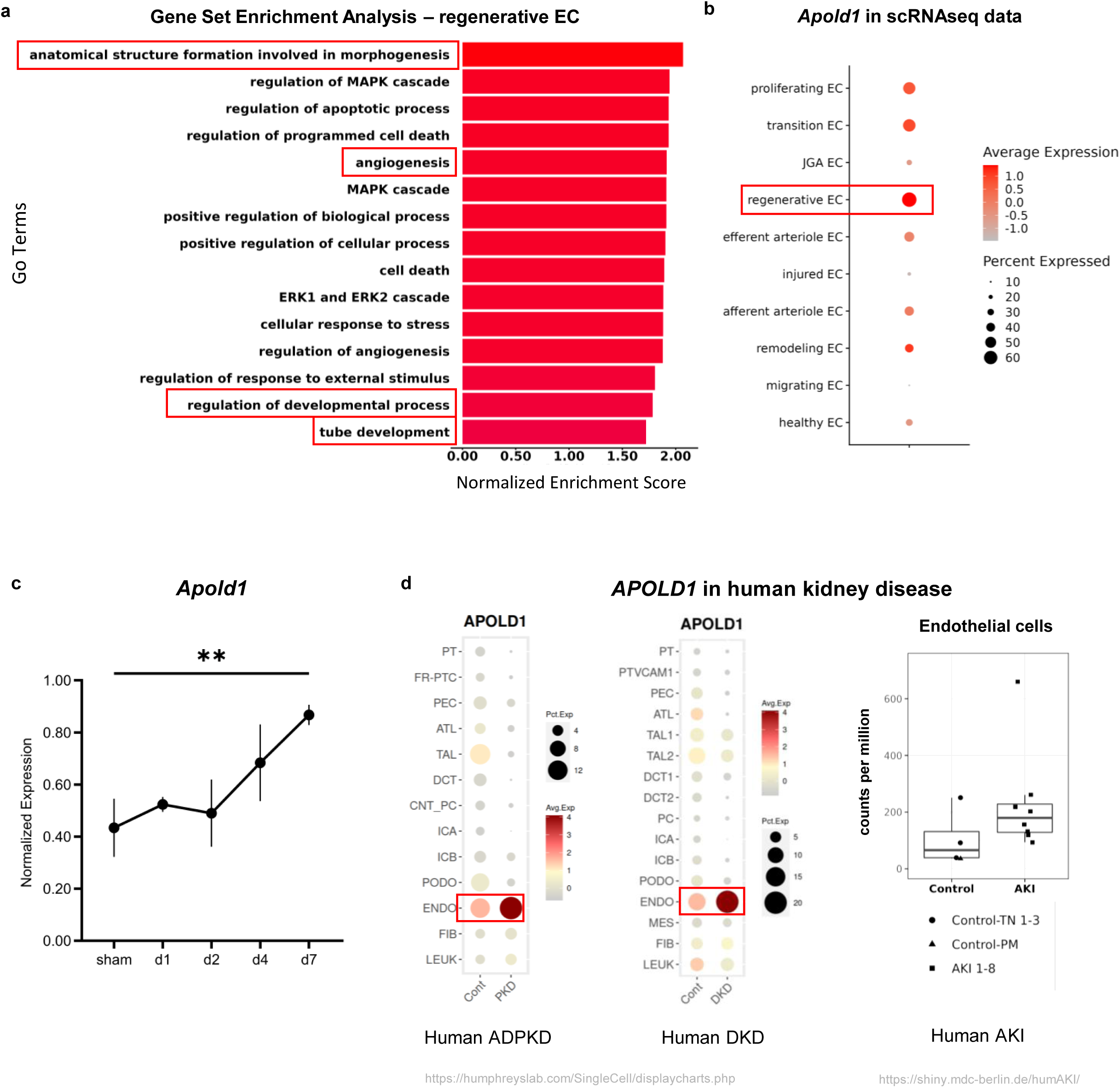
Single-cell transcriptomic profiling demonstrates injury-induced regulation of Apold1/APOLD1 in mice and humans. (a) Gene Set Enrichment Analysis of the regenerative endothelial cell (EC) cluster reveals the top enriched Gene Ontology (GO) terms (gene sets) associated with differentiation, angiogenesis, proliferation, and overall regeneration. Gene sets marked in red include *Apold1*. (b) Feature plot depicting the average expression of *Apold1* in every cluster of our single-cell RNA sequencing data. Up to 60% of cells in the regenerative EC cluster highly express *Apold1* (red box). (c) Normalized expression of *Apold1* throughout the experiment showing a significant increase in *Apold1* expression between sham and d7 post-injury in our single-cell RNA sequencing data. (d) *APOLD1* expression in single-cell RNA sequencing data from patients with autosomal dominant polycystic kidney disease (ADPKD), diabetic kidney disease (DKD), and acute kidney injury (AKI). Data taken from Wilson et al, 2022, Muto et al, 2022, Hinze et al, 2022. EC = endothelial cells; scRNAseq = Single-cell RNA sequencing

Notably*, Apold1* is highly upregulated in up to 60% of the cells associated with the *regenerative EC* cluster (Figure 5b). Normalized snRNAseq expression data demonstrated a significant increase in this gene during the regenerative phase (days 2–7) (Figure 5c). Using publicly available human kidney scRNAseq data, we examined *APOLD1* expression in patients with diabetic kidney disease (DKD)^59^, autosomal dominant polycystic kidney disease (ADPKD)^60^, and acute kidney injury (AKI)^61^. Across all datasets analyzed, *APOLD1* expression was consistently and significantly upregulated in patients with these diseases (Figure 5d), suggesting its potential role in EC activation and regeneration under pathological conditions in human kidneys as well. Analysis of publicly available human kidney snRNA sequencing data from patients with DKD^59^ and ADPKD^60^ revealed similar findings (Supplemental Figure 7).

### Apold1 deficiency impairs glomerular endothelial regeneration after injury

To assess the functional relevance of *Apold1* upregulation during regeneration, Apold1^-/-^ mice were subjected to the ECI model, and kidneys were harvested on days 1, 7, and 14 post-injury (Figure 6a).

**Figure 6.**
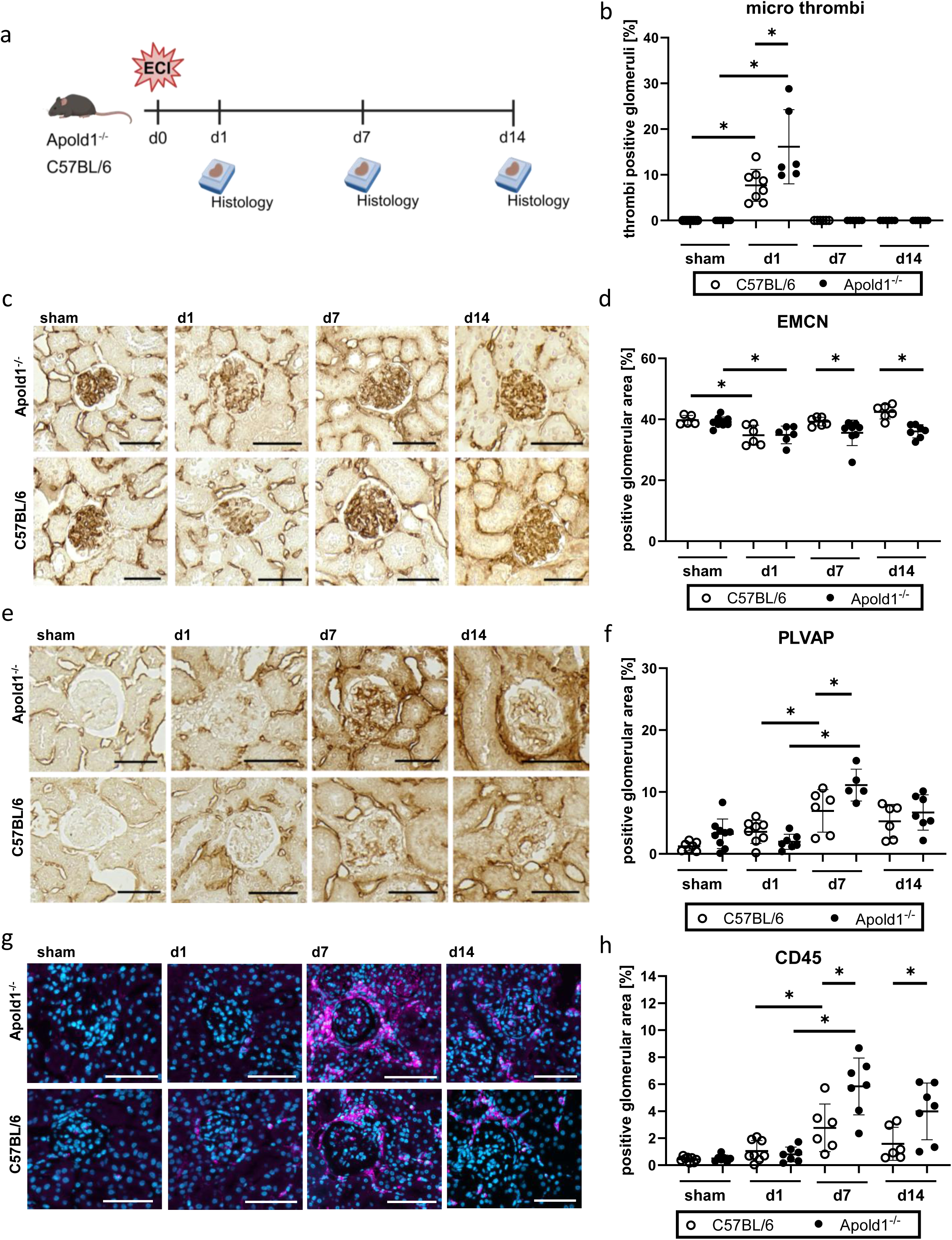
Apold1 deficiency impairs glomerular endothelial cell regeneration. (a) Experimental setup to investigate endothelial injury and regeneration in *Apold1^−/−^* mice. Kidney tissue was collected on day 1 (d1), day 7 (d7), and day 14 (d14) after induction of endothelial cell injury on day 0 (d0). Schematic created with BioRender. (b) Quantification of glomeruli containing thrombi in PAS-stained kidney sections from *Apold1^−/−^* and WT mice at d1, d7, and d14 post-injury. Data are shown as mean ± SD(c) Representative immunohistochemical staining for the endothelial cell marker endomucin in kidney sections from Apold1^−/−^ and WT mice. (d) Quantification of the endomucin-positive glomerular area in Apold1^−/−^ and WT mice at d1 and d7 post-injury. Sham-treated animals served as healthy controls. Data are shown as mean ± SD (e) Representative immunoperoxidase staining for PLVAP in kidney sections from Apold1^−/−^ and WT mice at d1, d7, and d14 post-injury. (f) Quantification of the mean PLVAP-positive glomerular area infrom Apold1^−/−^ and WT mice at d1, d7, and d14 post-injury. Sham-treated animals served as healthy controls. Data are shown as mean ± SD (g) Representative immunofluorescence staining for the immune cell marker CD45 in kidney sections from Apold1^−/−^ and WT mice at d1, d7, and d14 post-injury.. (h) Quantification of the CD45-positive cortical area in Apold1^−/−^ and WT mice at d1, d7, and d14 post-injury. Sham-treated animals served as healthy controls. Data are shown as mean ± SD.; *Apold1^-/-^* = *Apold1*-deficient mice, EMCN = endomucin; PLVAP= plasmalemma vesicle-associated protein,, =p<0.05. Scale bar = 50 µm.

PAS staining revealed significantly increased glomerular microthrombi formation in Apold1^-/-^ mice on day 1 compared with WT controls (Figure 6b). EMCN staining showed a significant reduction in EMCN-positive glomerular area in both genotypes on day 1, with no difference between Apold1^-/-^ and WT mice (34.9% vs. 34.8%; Figure 6c,d). While WT mice showed recovery towards sham levels by day 7 and day 14, Apold1^-/-^ mice displayed persistently reduced EMCN-positive areas at both time points, together with sustained abnormal endothelial morphology.

PLVAP staining, used as a marker of glomerular endothelial injury, increased in both genotypes on day 7 after injury. However, PLVAP-positive area was significantly higher in Apold1^-/-^ mice than in WT mice, indicating aggravated endothelial injury on day 7 (Figure 6e,f). At day 14, PLVAP expression remained elevated in both WT and Apold1-/- mice compared with sham-operated animals. However, no significant difference was detected between the genotypes. In addition, CD45-positive leukocyte infiltration was significantly increased in Apold1^-/-^ mice on days 7 and 14 compared with WT controls, suggesting enhanced inflammatory activity (Figure 6g,h).

## DISCUSSION

This study provides a comprehensive characterization of gECs and their subpopulations during regeneration after EC-specific renal injury using scRNAseq and ST. Although previous studies suggested a role for intrinsic renal ECs in repair, the precise cellular origin of regenerated gECs remained incompletely defined ^62^. Using Cdh5(PAC)-ERT2-tdT lineage tracing, we showed that EC regeneration after ECI mainly originates from pre-existing adult renal ECs rather than non-endothelial precursor cells.

Based on this finding, we performed scRNAseq across several time points after ECI, which revealed 10 gEC clusters with distinct temporal patterns and potential functions, mapping the progression from injury to near-complete structural recovery. Xenium ST localized these subpopulations within the glomerulus and confirmed their heterogeneous spatial distribution. These data are consistent with the idea that repair-associated endothelial states are broadly distributed rather than confined to a single spatially restricted EC pool^63,64^.

Using ST, we were able to address concerns about preparation-induced artifacts in scRNAseq data, as highlighted by Van den Brink and colleagues^65^, by showing high concordance between our scRNAseq and ST data. This argues against the possibility that the high abundance of IEGs in the regenerative EC cluster is solely caused by dissociation artifacts and supports a role for local stress-response programs in endothelial regeneration. Thus, although dissociation-associated induction of immediate early genes remains an important caveat in single-cell studies, their spatial confirmation in tissue supports the interpretation that at least part of this program reflects genuine in situ endothelial activation during repair.

The temporal organization of gEC states suggests that endothelial repair follows a phased but overlapping trajectory, beginning with inflammatory stress adaptation and progressing toward migratory, proliferative, and remodeling programs. Rather than representing strictly sequential stages, these phases likely reflect co-existing endothelial responses within the injured glomerulus, where some cells undergo loss or dysfunction while others activate survival and repair-associated pathways (Figure 7). High Serpina3 expression in the injured EC cluster suggests activation of survival-associated pathways, including PI3K/AKT and MAPK/ERK1/2 signaling^66^. In parallel, a transition EC state emerged that combined inflammatory, migratory, and regeneration-associated programs^34,36^ potentially linking injury-induced inflammation^67^ with pro-regenerative remodeling^68^ and cross-talk with neighboring glomerular cells ^32,33,69^. The increased abundance of migrating ECs on day 2 further supports active endothelial migration toward injured vascular areas^7^. This transitional state may represent a key decision point at which injured endothelial cells either progress toward repair or remain in a dysfunctional inflammatory state. Future ligand–receptor analyses could help determine whether podocyte- or mesangial-derived signals contribute to this endothelial state transition.

**Figure 7.**
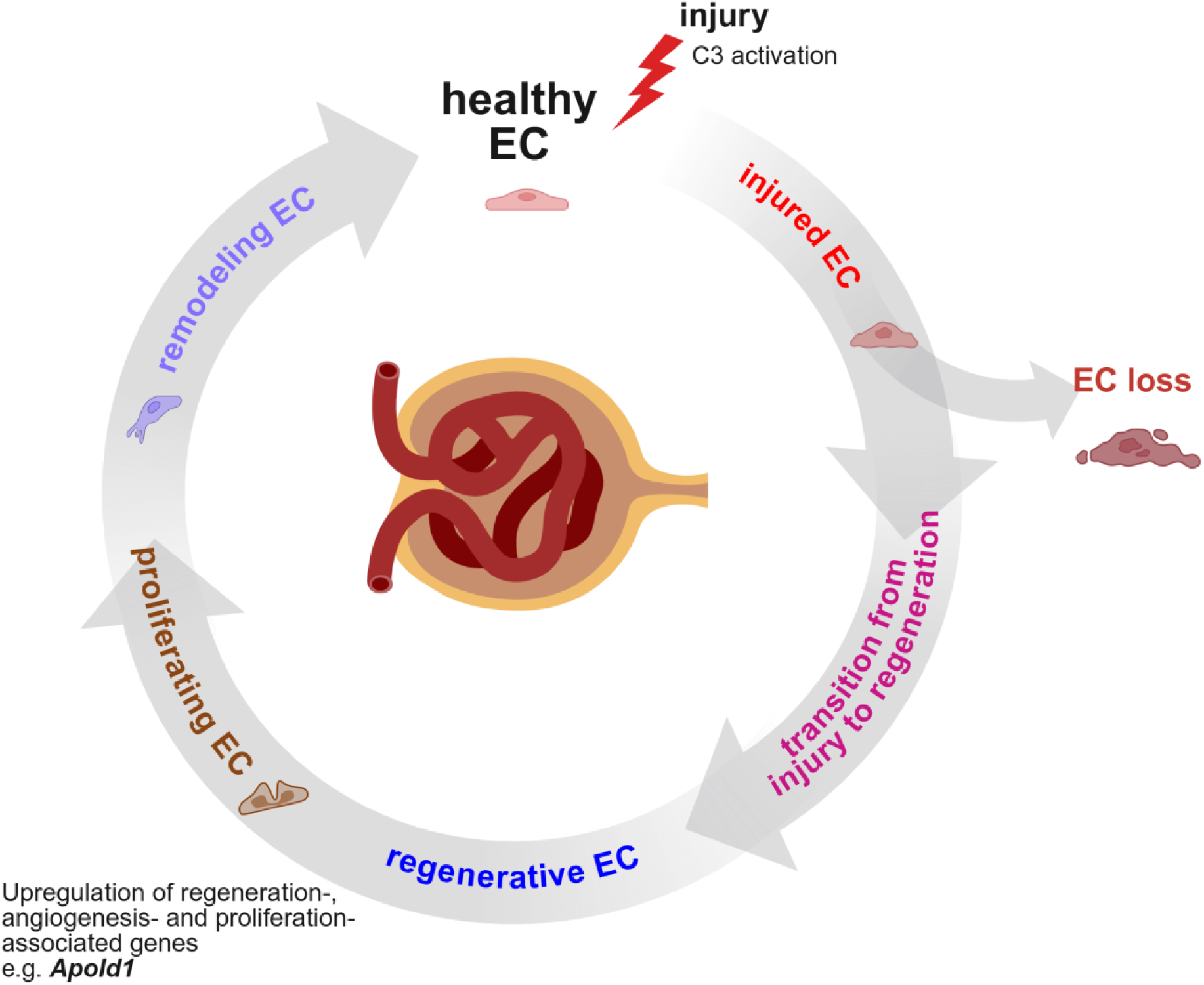
Potential mechanism for endothelial cell regeneration. Potential process of renal endothelial injury and the subsequent regenerative phases, as suggested by our single-cell RNA sequencing and spatial transcriptomics data. The differently colored endothelial cell cluster represent the various transcriptional patterns found during the injury and regeneration phase. Image created in BioRender. EC = endothelial cell, scRNAseq = single-cell RNA sequencing; ST = spatial transcriptomics

Between days 4 and 7, regenerative, remodeling, and proliferating EC states became dominant. In the *regenerative EC* cluster, the MAPK/JNK signaling pathway is highly enriched, which is known to enhance *VEGF* production^70,71^, a key driver of EC proliferation and migration^72^. Previous studies from our group have already evaluated VEGF expression in similar models, confirming its essential role in renal vascular repair^5,73^. The prominent expression of IEGs in the regenerative EC subpopulation may indicate an acute activation program associated with endothelial stress adaptation, proliferation, and migration. This could mark a significant shift towards the restoration of vascular function^74^. For example, *Atf3*, an IEG, has been experimentally identified as a key marker in EC subpopulations with regenerative potential, as demonstrated in large vessels and lung tissue^49,75,76^. Within this group, the EC-specific gene *Apold1* emerged as an interesting candidate for functional validation, given its endothelial-restricted expression and its established role in pathological angiogenesis^13^.

By day 7 after ECI, *proliferating* and *remodeling* EC clusters were strongly represented. These clusters expressed genes linked to cell division (*Mki67*^56^ and *Cenpf*^55^), vascular growth (*Hexim*^46,77^), endothelial stability (*Adamts1*^43^ and *Btg2*^47^), stress adaptation (*Jun*^78^ and *Fos*^79^), and anti-fibrotic responses (*Plk2*^48,80^), suggesting that late repair involves both EC expansion and structural stabilization of the regenerated endothelium^44,81^

The human biopsy data suggest that selected endothelial injury and repair programs identified in the mouse ECI model are also detectable in human glomerular disease. However, the differences between TMA subtypes indicate that these programs are modulated by the underlying pathogenic trigger rather than representing a uniform endothelial response. The observation that similar endothelial injury-associated signatures are present across TMA, DKD, and ADPKD suggests that part of the glomerular endothelial response may represent a conserved endothelial injury response shared across different kidney diseases rather than a disease-specific signature. Nevertheless, the relative contribution of individual pathways likely depends on the dominant pathogenic trigger, such as metabolic stress, cyst-associated remodeling, complement activation, or hemodynamic injury.

Further analysis of publicly available human scRNA-seq datasets from patients with DKD, ADPKD, and kidney allograft rejection revealed increased APOLD1 expression in endothelial cells across all three disease settings^59–61^. This consistent upregulation across independent datasets suggests that *APOLD1* may be part of a broader endothelial response to renal injury and disease rather than being restricted to one specific condition. Together with previous reports showing *APOLD1* induction under metabolic stress, including hypoxia and ischemia^82–84^ these findings support its potential clinical relevance and identify *Apold1* as a candidate regulator of vascular repair warranting further investigation in experimental models of renal injury and regeneration.

Our animal experiments in *Apold1*-deficient mice revealed increased glomerular microthrombus formation after renal injury, supporting the prothrombotic phenotype associated with *Apold1* deficiency^85^. Notably, although the extent of initial endothelial damage was comparable to WT mice, Apold1^−/−^ mice subsequently showed impaired endothelial regeneration as indicated by endomucin staining, increased endothelial dysfunction^84^ as indicated by PLVAP expression, and persistently increased leukocyte infiltration. These findings indicate that *Apold1* is not required to prevent the acute injury itself, but is important for endothelial repair, similar to its function in other organs such as brain and skeletal muscle^13,86^. They also support a role for *Apold1* as an endothelial-intrinsic regulator of glomerular vascular repair.

Consistent with this repair-associated state, the regenerative endothelial cluster showed enrichment of MAPK/ERK signaling, suggesting that *Apold1* may be linked to injury-induced pathways that coordinate endothelial activation, angiogenesis, and vascular remodeling. This interpretation is consistent with the known role of MAPK/ERK signaling in regeneration, angiogenesis, endothelial migration, and vascular remodeling^87^. Based on reported functions of *Apold1* in endothelial cell permeability, cell-cell contacts, and angiogenesis, a delayed *Apold1* response may support vascular remodeling and the restoration of vascular function.

The combination of impaired EMCN recovery, increased PLVAP expression^36,88^, and persistent inflammation in Apold1^−/−^ mice suggests that Apold1 deficiency may keep endothelial cells in an activated, immature, and/or dysfunctional state. Together, these findings support a model in which Apold1 acts as an injury-induced endothelial regulator that is dispensable for baseline vascular homeostasis but required for efficient repair under pathological conditions. In the kidney*, Apold1* may promote glomerular endothelial regeneration by facilitating endothelial re-differentiation, limiting persistent endothelial dysfunction, and supporting the resolution of injury-associated inflammation.

Despite these promising findings, our study has limitations that warrant consideration. Although the restoration of the EMCN-positive glomerular endothelial area indicated that structural endothelial repair was largely completed by day 14 after ECI, our time course does not allow definitive conclusions regarding complete functional healing or long-term transcriptional normalization of regenerated gECs. Our investigation primarily focused on gECs. The renal microvasculature is highly heterogeneous, and it is possible that peritubular capillary ECs undergo distinct injury responses and regenerative pathways that differ from the process described here. Although the combination of scRNAseq and ST allowed us to temporally and spatially map EC subpopulations, these techniques inherently capture static “snapshots” of cellular states. For the ST data, segmentation limitations remain, especially within the glomerular area, where the close proximity of various cell types hinders precise cell differentiation and limits the data to a near single-cell resolution. Additionally, compared to the Visium platform from 10x Genomics, which can profile a larger number of genes, our Xenium approach was limited to a predefined panel of 400 genes. While we can infer the transition of ECs from the inflammatory phase to the remodeling phase, continuous *in vivo* longitudinal tracking of a single cell’s fate, especially with such a complex gene expression pattern as our subpopulations, is still technologically constrained. Finally, while we successfully identified *Apold1* as a candidate endothelial-intrinsic regulator of glomerular vascular repair, the precise downstream molecular targets and its potential cross-talk with neighboring glomerular cells, such as podocytes and mesangial cells, require further detailed functional investigation in the future.

In conclusion, we identified and localized previously unknown gEC subpopulations and states involved in injury and regeneration during ECI. The temporal pattern of gEC regeneration suggests a dynamic, multi-phased process driven by local signaling and environmental factors encompassing the ECI response, EC migration, and remodeling, as well as EC proliferation to recover the endothelium (Figure 7). We observed the upregulation of the gene *Apold1* during the regenerative phase of the process and identified it as a new potential regulator of glomerular vascular repair. Its disease-associated regulation in human kidney disease datasets suggests potential clinical relevance and supports further investigation of APOLD1 as a target to enhance renal microvascular repair. To our knowledge, this study provides the first temporally and spatially resolved map of glomerular endothelial regeneration after selective endothelial injury.

## DATA SHARING STATEMENT

The scRNAseq and ST data supporting the findings of this study are openly available in Gene Expression Omnibus (GEO) under accession numbers GSE285062 and GSE284472.

## AUTHOR CONTRIBUTIONS

Conceptualization: J.S.; C.H.; M.S., Formal analysis: M.S.; J.S, Funding acquisition: C.H.; J.S.; V.T., Investigation: M.S.; J.S.; H.W; P.A.; L.D; P.H; A.W.; Ma.S.; T.C.; J.A., Methodology: J.S.; T.C.; J.A., Project administration: J.S., Software; M.S., Supervision: J.S.; C.H., Writing – original draft: M.S.; J.S.; C.H, Writing – review& editing: C.H.; V.T.; J.S.; M.S.

## DECLARATION OF GENERATIVE AI AND AI-ASSISTED TECHNOLOGIES IN THE WRITING PROCESS

During the preparation of this work, the authors used Gemini Pro 3.1. in order to clarify and improve readability of the human-written text. After using this tool, the authors reviewed and edited the content as needed and take full responsibility for the content of the publication.

## SUPPLEMENTARY MATERIAL

**Supplementary Methods and Supplementary References.**

**Supplementary Figure S1** [Lineage-tracing of regenerating endothelial cells]

**Supplementary Figure S2** [Experimental setups for animal experiments and transcriptomic analyses]

**Supplementary Figure S3** [Quality control analyses of single-cell RNA sequencing data]

**Supplementary Figure S4** [Single-cell RNA sequencing reveals new glomerular endothelial cell populations during endothelial cell injury compared to healthy animals]

**Supplementary Figure S5** [Comparability of gene expression of single-cell RNA sequencing and spatial transcriptomics data]]

**Supplementary Figure S6** [Selected scRNAseq markers are present in human kidney samples of complement-mediated TMA]

**Supplementary Figure S7** [Expression pattern of selected markers based on databases]

**Supplementary Table S1** [List of antibodies used for flow cytometry and immunohistochemistry]

**Supplementary Table S2** [Top 50 DEGs of single-cell RNA sequencing data]

**Supplementary Table S3** [Gene Set Enrichment Analysis of the regenerative EC cluster]

**Supplementary Table S4** [Marker genes for spatial transcriptomics]

**Supplementary Table S5** [ Clinical renal function parameters in patients]

**Supplemental Methods**

## ACKNOWLEDGMENTS

This work was funded by the Deutsche Forschungsgemeinschaft (DFG, German Research Foundation) grant number HU 600/13-1. We are grateful to Prof. Dr. Peter Boor, Ph.D. for providing the Cdh5(PAC)- CreERT2 mice. We are also grateful to Johannes Bohacek and Katrien De Bock from ETH Zurich for kindly providing the Apold1-deficient mice. We thank Dr. Anne Gompf and Katja Bernhardt from the CMCB Flow Cytometry facility for sorting assistance. We would like to acknowled Susanne Reinhardt, Dr. Fabian Rost, Dr. Andreas Petzold, and the whole single-cell unit of the DRESDEN-concept Genome Center for their advice and support in single-cell RNA sequencing. We thank Izabela Plumbom and the team from the BIH/MDC Genomics Technology Platform in Berlin for conducting spatial transcriptomics (Xenium). We also thank Silke Tulok and the team of the MTZ – Core Facility Cellular Imaging at the medical faculty Carl Gustav Carus for imaging support. Finally, we would like to thank the entire Experimental Nephrology research group for their support during the experiments and throughout the writing process of this paper.

