## Supplementary Figures captions for "Spatiotemporal Mapping of Glomerular Endothelial Regeneration Identifies *Apold1* as an Intrinsic Regulator of Repair"

### Supplementary Figure S1

#### Lineage-tracing of regenerating endothelial cells

(a) Experimental setup for lineage tracing of endothelial cells after injury in *Cdh5(PAC)-ERT2-tdTomato* reporter mice. After 5 days of tamoxifen induction and a wash-out phase, a partial biopsy was done on the left kidney 24h before induction (-d1) of the endothelial cell injury model (d0). A second partial biopsy was done 24 h after injury (d1). Kidneys were harvested on day seven. Kidney samples were used for flow cytometry and histological analyses. Image created in BioRender (b) Number of *ERG*-positive nuclei per glomerulus shows a significant decrease at day 1 compared to day 7 and sham mice. (c) Analysis of tdTomato- and ERG-positive cells of sham-operated and ECI mice showing a mean tdTomato-positive/*ERG*-positive proportion of over 85 %. (d) Representative pictures of mouse kidney glomeruli stained against the endothelial nuclear marker ERG, DAPI, and tdTomato at day one and day seven after injury, as well as control (sham-operated mice). Data are represented asmean ± SD; scale bar = 10 µm, EC = Endothelial Cell; ECI = Endothelial Cell Injury, ERG = ETS Transcription Factor ERG

### Supplementary Figure S2

#### Experimental setups for animal experiments and transcriptomic analyses

(a) The endothelial cell injury (ECI) model mimics a human thrombotic microangiopathy and induces kidney-specific endothelial cell damage. Concanavalin-A is injected via the left renal artery and binds specifically to the endothelium, followed by an anti-concanavalin A serum produced in rabbit. This leads to the activation of the mice's complement system which eventually damages the endothelial cells. Kidney tissue is taken at earliest 24 hours after the model induction. Image created in BioRender (b) For the single-cell sequencing approach, glomerular endothelial cells are isolated from fresh kidneys of three mice in each replicate, and staining is performed using oligo-antibodies (Hashtag AB) to trace the origin of the cells. Cells are sorted by FACS to make sure that only the desired cell type is used further. After library preparation, cells are sequenced. Image created in BioRender. (c) Three kidneys are taken at each time point for Spatial Transcriptomics analysis. The kidneys are fixed in a PFA solution, embedded in paraffin, cut and analysed on the 10x Genomics Xenium platform.. The libraries used were based on the 10x-provided mouse-tissue panel and a custom-made gene panel with genes selected based on our single cell RNA sequencing data. Image created in BioRender. (d) Tables showing the number of animals used per mouse strain and experimental approach. ECI = endothelial cell injury; FACS = fluorescence-activated cell sorting; PFA = paraformaldehyde

### Supplementary Figure S3~~2~~

#### Quality control analyses of single-cell RNA sequencing data

(a) Uniform Manifold Approximation and Projection (UMAP) plot visualizing endothelial cells isolated from glomeruli and their expression of endothelial cell marker *Cdh5* and *Ehd3*~~.~~ ~~–~~(b) To evaluate the presence of potential contaminating cells in the single-cell RNA sequencing data, the data was filtered for marker of common renal cell types. The UMAP plots are color-coded to highlight expression of specific markers, such as *WT1*, *Nphs2* for podocytes; *Itga8* and *Des* for mesangial cells, *Hbb-bt* and *Hbb-bs* for red blood cells. Cells with these markers were excluded from subsequent analyses~~;~~. ~~Uniform Manifold Approximation and Projection~~

### Supplementary Figure S4

#### Single-cell RNA sequencing reveals new glomerular endothelial cell populations during endothelial cell injury compared to healthy animals

(a) Scaled heatmap with averaged expression of single-cell RNA sequencing data of the top 20 differentially expressed genes (DEG) for each cluster of glomerular endothelial cells from healthy sham-operated mice. (b) Cluster analyses of glomerular endothelial cells separate per time point. Cluster colors and numbers are randomly associated for every time point. (c) Principal Component Analysis (PCA) plot depicting the Jaccard similarity coefficient of glomerular endothelial cell subpopulations in endothelial cell injury mice. The cluster projection illustrates the transformation of control glomerular endothelial cell phenotypes from endothelial cells after injury on days 1, 2, 4, and 7. '???' denotes unknown or novel clusters. (d) Uniform Manifold Approximation and Projection (UMAP) plot of control and injured endothelial cells after collective cluster analysis. Cells are colored according to their time point collected; ECI = endothelial cell injury; gEC = glomerular endothelial cells;

**Supplementary Figure S5**

**Comparability of gene expression of single-cell RNA sequencing and spatial transcriptomics data**

(a) Transcript density maps of markers *C3*, *Bst2* and *Egr1* throughout the experiment. scale bar = 1000 –µm (b) Changes in normalized expression of markers *C3*, *Bst2* and *Egr1* throughout the experiment. *C3* is upregulated on day 1 (d1) as a result of the endothelial cell injury induction. *Bst2* is upregulated on d1, especially in injured endothelial cells. The gene Egr1 increases in the later phase of the process during regeneration (day ?). (c) Normalized expression of one of the top markers from every cluster, compared to the normalized expression of the same marker in spatial transcriptomics, to show comparability of the data. scRNAseq = single-cell RNA sequencing; ST = spatial transcriptomics

**Supplementary Figure S6**

**Selected scRNAseq markers are present in human kidney samples**

Histological stainings of important markers found in scRNAseq data. Tissue was taken from patients (Supplementary Table S6) with Minimal Change Glomerulus Neprithis (MCGN), complement-mediated thrombotic microangiopathy (TMA) and malignant hypertensive TMA. Brown color shows the presence of proteins Serpina3, Atf3, DNAJB1, Egr1, BST2 and Sox17 which are all associated with either injury or regeneration in our TMA-like mouse model. NC = negative control; MCGN=minimal change glomerular nephritis; TMA = thrombotic microangiopathy; scale bar = 50 µm

**Supplementary Figure S7**

**Expression pattern of selected markers based on databases**

To show the expression of selected markers, published scRNA-seq and snRNA-seq datasets of kidney cells from healthy controls and from patients with diabetic kidney disease (DKD) or autosomal dominant polycystic kidney disease ADPKDwere analyzed. The expression of SERPINA3, ATF3, DNAJB1, EGR1, BST2, and SOX17 is shown. Endothelial cell populations are marked with red boxes.
