## Supplementary Figures for "Spatiotemporal Mapping of Glomerular Endothelial Regeneration Identifies *Apold1* as an Intrinsic Regulator of Repair"

a

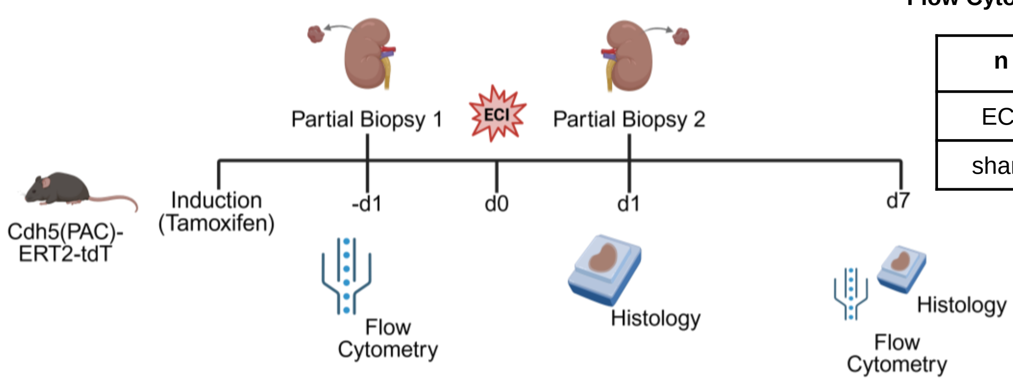


**Flow Cytometry** (Cdh5(PAC)-ERT2-tdTomato)

| n | -d1 | d1 | d7 | total |
|---|---|---|---|---|
| ECI | 12 | 7 | 12 | 35 |
| sham | 7 | - | 7 | |

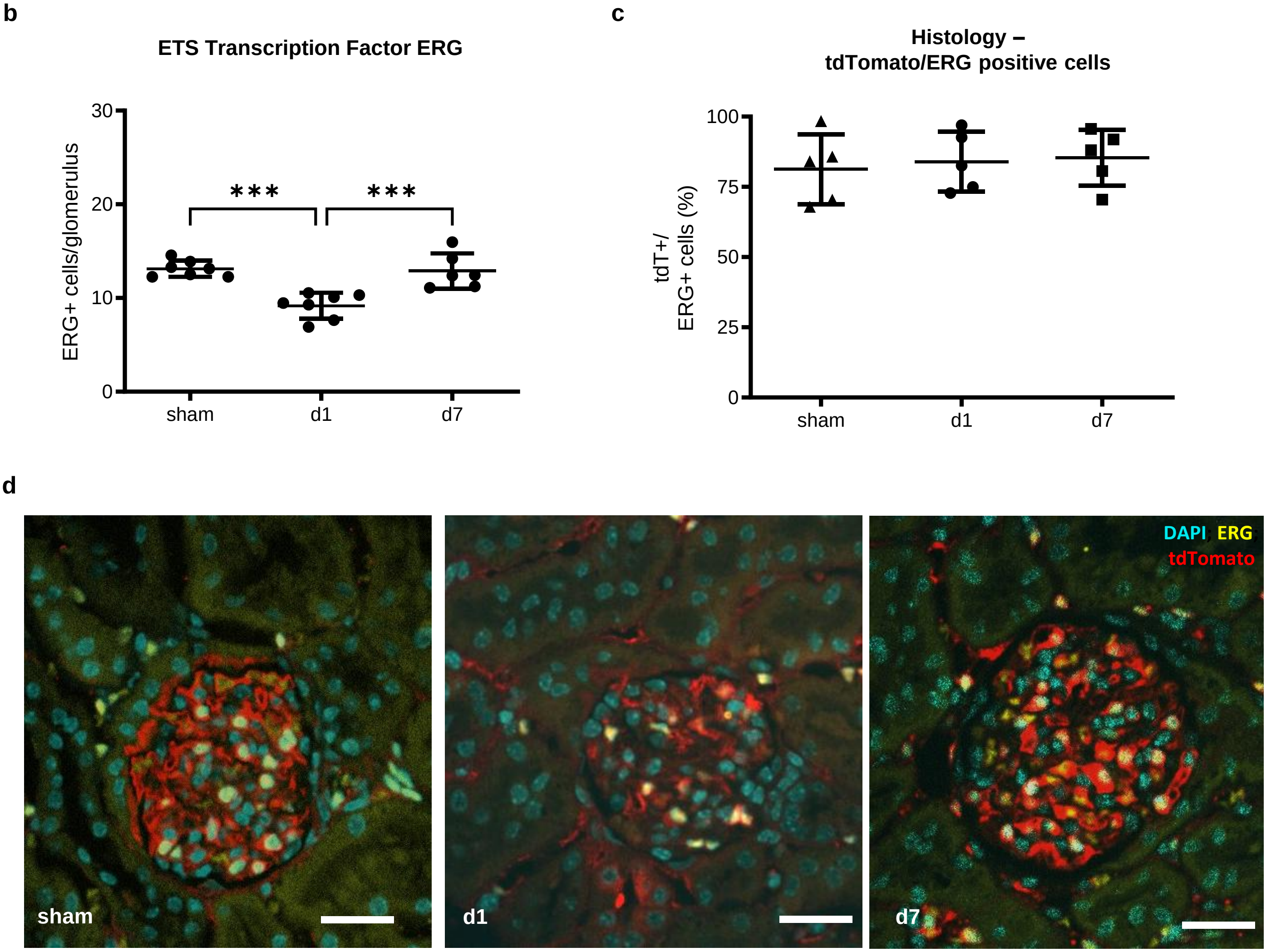


**Supplementary Figure S1**

**a**

**scRNAseq** (Tg(TIE2GFP)287Sato/J) **& ST (**C57BL/6JRj)

| n | sham | d1 | d2 | d4 | d7 | total |
|---|---|---|---|---|---|---|
| Tg(TIE2GFP) 287Sato/J | 9 | 9 | 9 | 9 | 9 | 45 |
| C57BL/6JRj | 2 | 3 | 3 | 3 | 3 | 14 |

**Single-cell RNA sequencing**

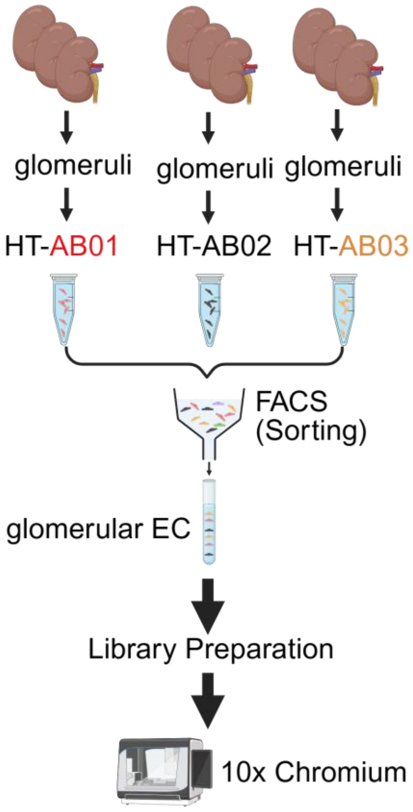


**b** **Spatial Transcriptomics**

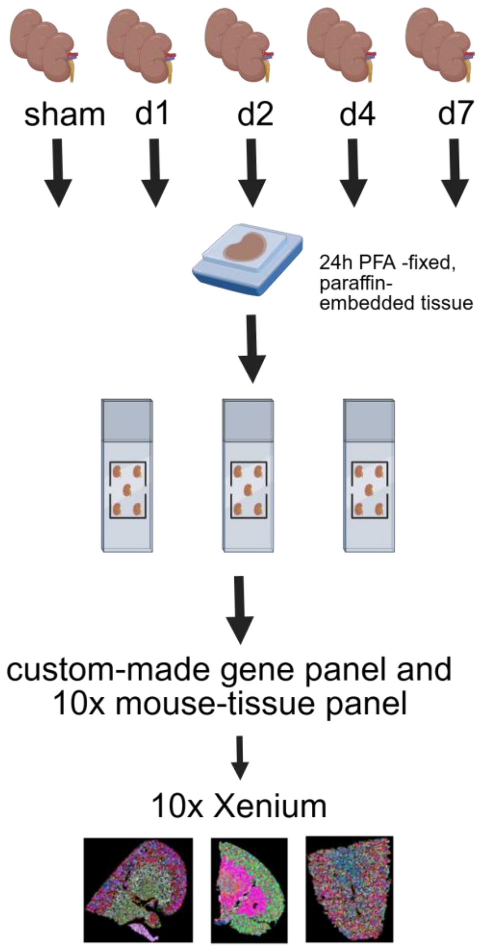


**c** **Apold1-/-**

| n | sham | d1 | d7 | d14 | tot al |
|---|---|---|---|---|---|
| C57BL/6 J-Apold1tm1(LacZ) | 9 | 6 | 8 | 7 | 30 |
| C57BL/6JRj (WT) | 5 | 6 | 6 | 6 | 23 |

**Supplementary Figure S2**

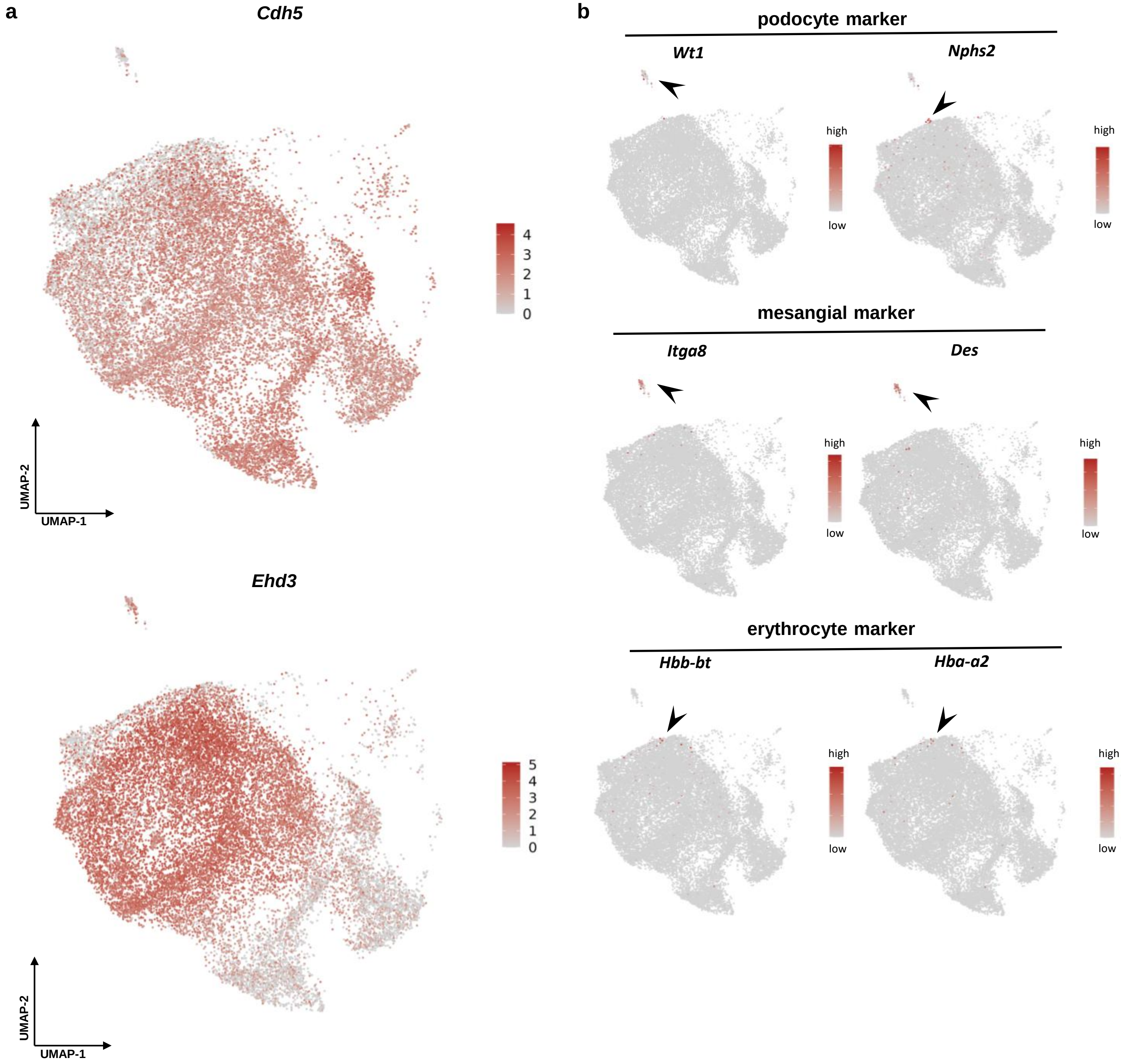


Supplementary Figure S3

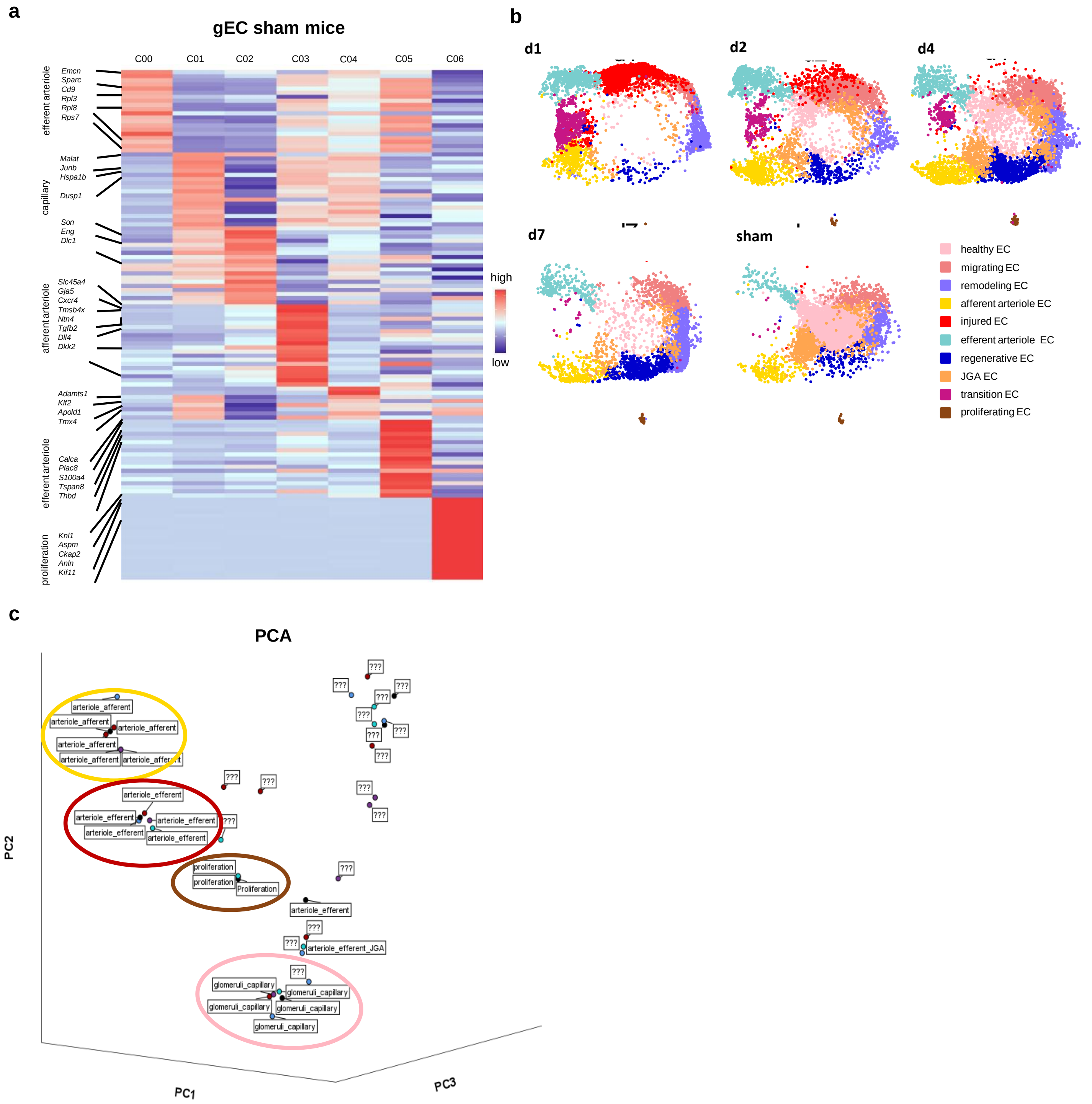


**Supplementary Figure S4**

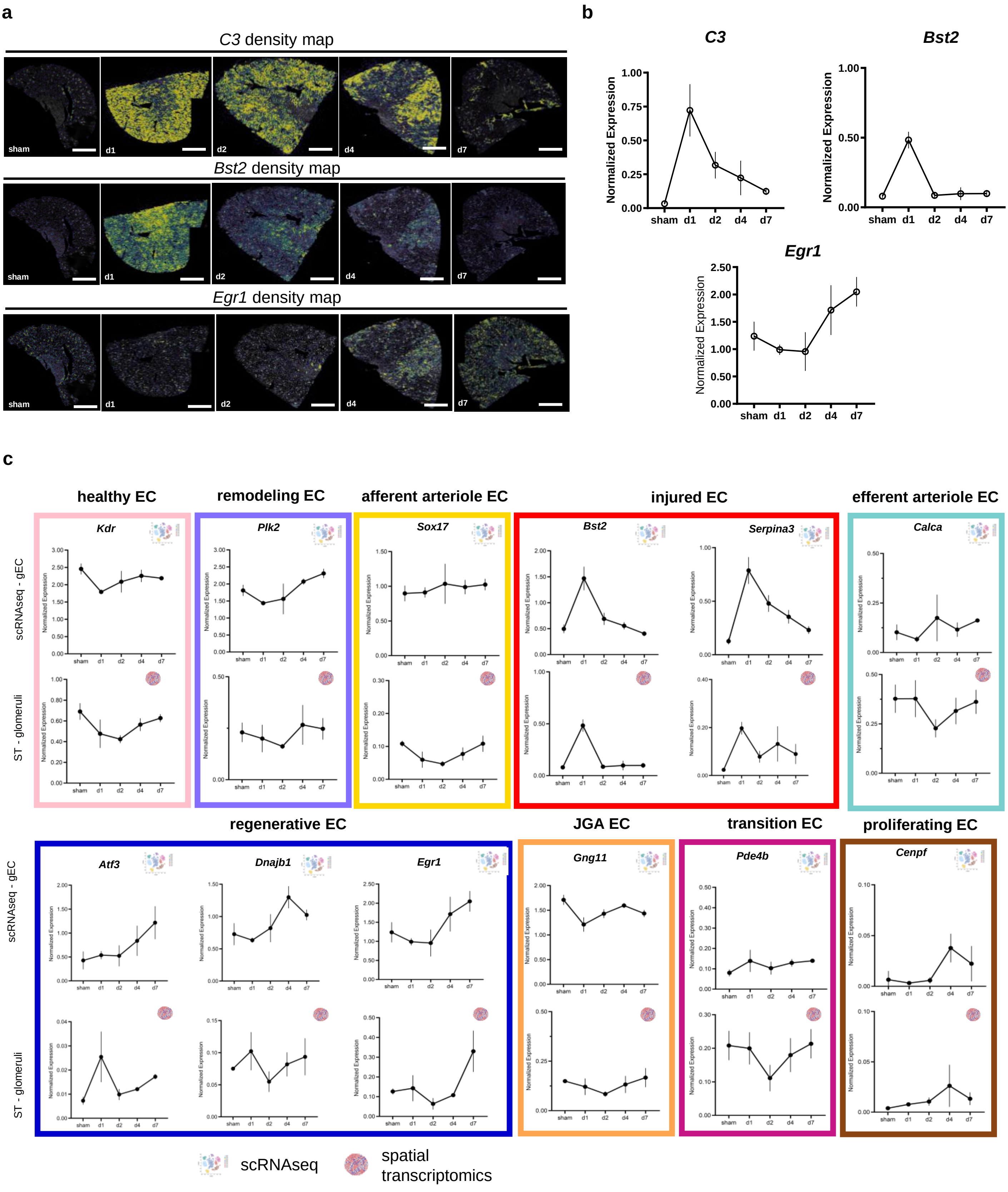


Supplementary Figure S5

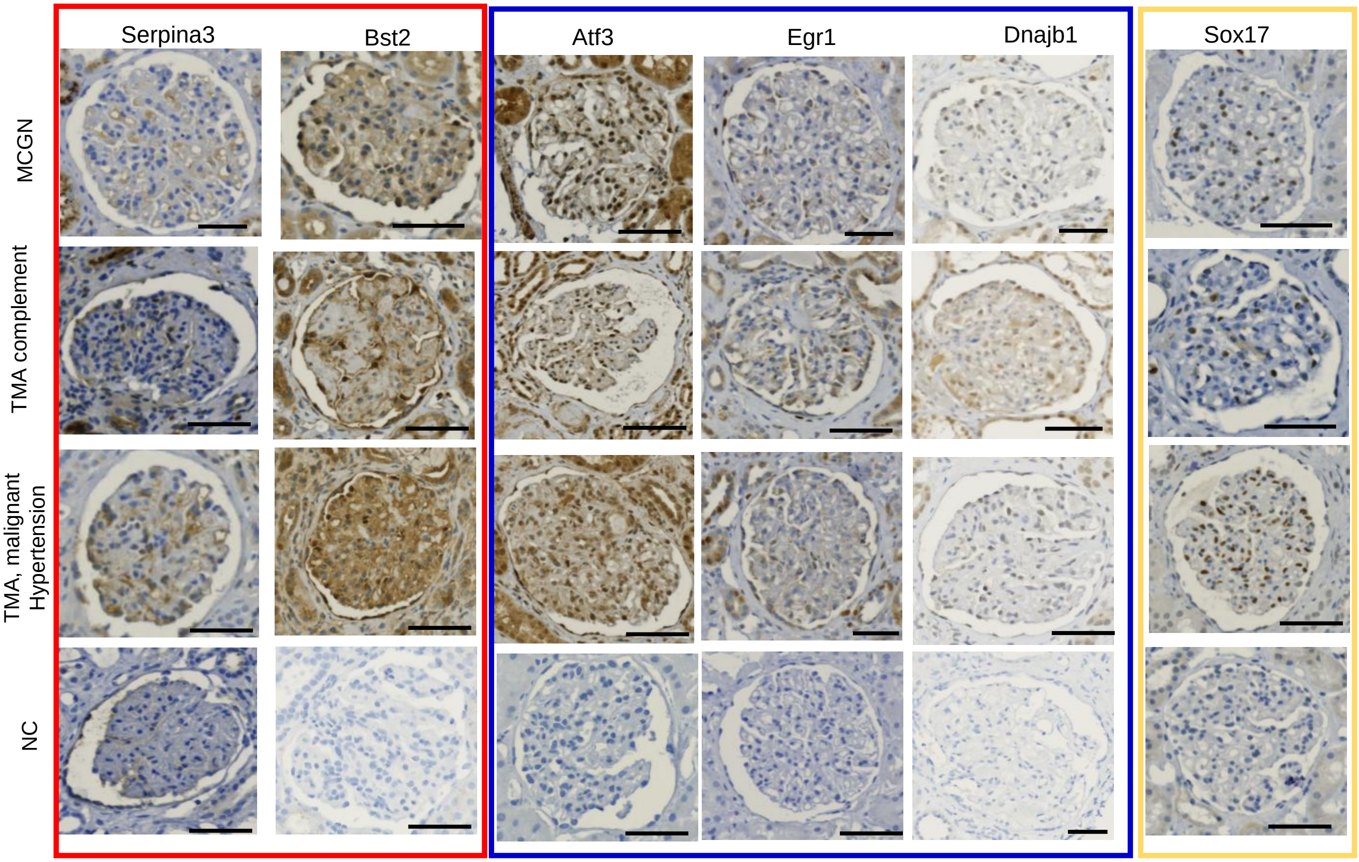


**Supplementary Figure S6**

Human ADPKD snRNA + scATAC-seq: [Muto et al, Nat Commun 2022](#).

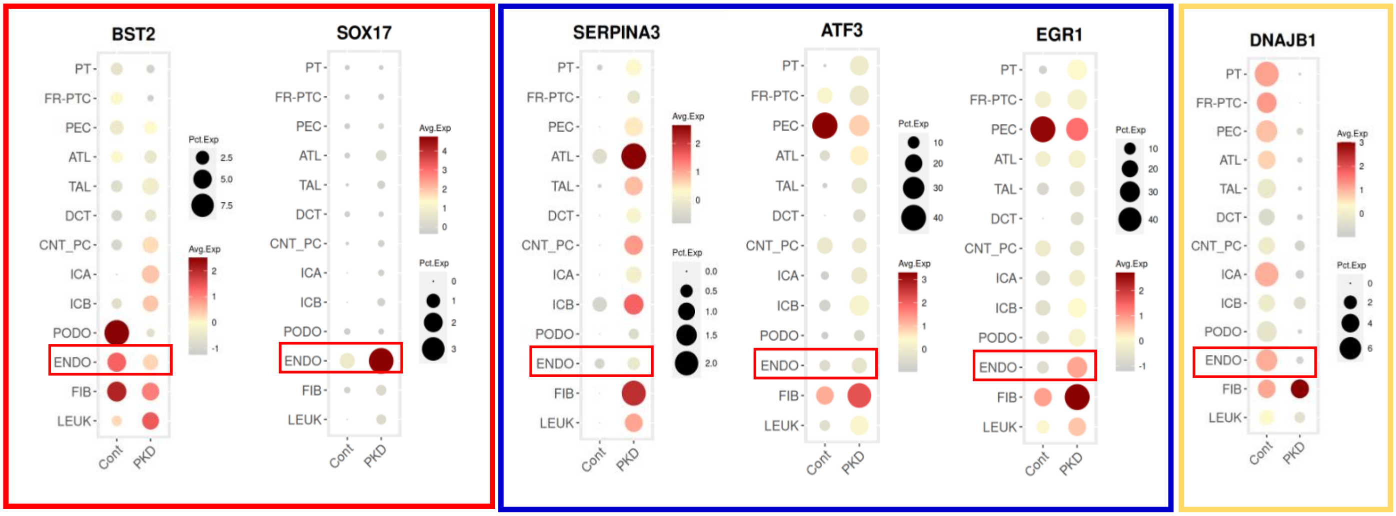


Human DKD snRNA + scATAC-seq: [Wilson and Muto et al. Nat Commun 2022](#).

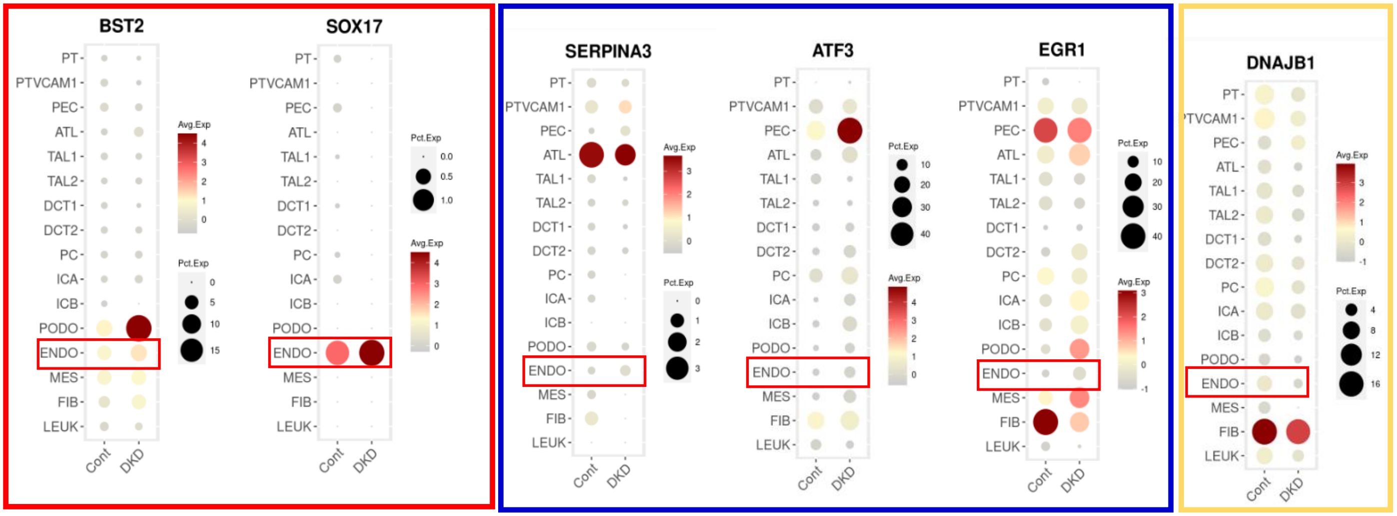


**Supplementary Figure S7**
