## Supplementary tables for "Spatiotemporal Mapping of Glomerular Endothelial Regeneration Identifies *Apold1* as an Intrinsic Regulator of Repair"

| ANTIBODIES | SOURCE | IDENTIFIER |
|---|---|---|
| Rat anti-mouse CD31 FITC (clone 390) | Thermo Fisher Scientific | Cat#11-0311-82RRID: AB_465012 |
| Rat anti-mouse CD45 PE-Cy7 (clone 30-F11) | Thermo Fisher Scientific | Cat#25-0451-82RRID: AB_2734986 |
| Rat anti-mouse CD105 eFlour™ 450 (clone MJ7/18) | Thermo Fischer Scientific | Cat# 48-1051-82 RRID:AB_10548959 |
| Rat anti-mouse CD73 PerCP/Cyanine5.5 | BioLegend | Cat#127213 RRID: AB_11219608 |
| Rat anti-mouse ICAM-2 Alexa Fluor 647 | Thermo Fisher Scientific | Cat#A15452 RRID: AB_2534465 |
| Rabbit anti-mouse IgG2 tdTomato antibody (clone 16D7) | Kerafast | Cat# EST203 RRID:AB_2732803 |
| Rabbit Alexa Fluor® 647 Anti-ERG antibody (clone EPR3864) | abcam | N/A |
| Rat anti-mouse IgG2a Endomucin antibody (clone V.7C7) | Santa Cruz Biotechnology | Cat# sc-65495 RRID: AB_2100037 |
| Rabbit anti-human SERPINA3 Antibody (polyclonal) | Atlas Antibodies | Cat# HPA000893 RRID:AB_10794315 |
| Rabbit anti-human EGR1 Antibody (clone 15F7) | Cell Signaling Technology | Cat# 4153 RRID:AB_2097038 |
| Rabbit anti-human ATF3 Antibody (polyclonal) | Atlas Antibodies | Cat# HPA001562 RRID:AB_1078233 |
| Rabbit anti-human SOX17 (polyclonal) | R&D Systems | Cat# AF1924 RRID:AB_355060 |
| Goat Anti-Rat IgG Antibody (H+L), Biotinylated | Vector Laboratories | Cat# BA-9400 RRID: AB_2336202 |
| Goat Anti-Rabbit IgG Antibody (H+L), Biotinylated | Vector Laboratories | Cat# BA-1000, RRID:AB_2313606 |
| Rabbit Anti-concanavalin A serum | Sequence Laboratories GmbH | N/A |
| Rabbit Anti-BST2 (polyclonal) | Merck (Sigma-Aldrich) | HPA017060 |
| Rabbit Anti-DNAJB1 (ployclonal) | Merck (Sigma-Aldrich) | HPA063247 |

### Supplementary Table S2: Top 50 DEGs of single-cell RNA sequencing data

A list of the top 50 differentially expressed genes of the 10 glomerular endothelial cell clusters identified by single-cell RNA sequencing.

| Rank | healthy EC | migrating EC | remodeling EC | afferent arteriole EC | injured EC | efferent arteriole EC | regenerative EC | JGA EC | transition EC | proliferating EC |
|---|---|---|---|---|---|---|---|---|---|---|
| 1 | Igfbp5 | Syne1 | Pcdh17 | Vim | Ly6a | Tspan8 | Atf3 | Ntn4 | Igfbp3 | Cenpf |
| 2 | Sapcd1 | Lars2 | Plk2 | Sat1 | Serpina3g | S100a4 | Egr1 | Slc45a4 | Plvap | Hmgb2 |
| 3 | Gchfr | Son | Hexim1 | Edn1 | Cd74 | Calca | Dnajb1 | Plat | Serpine1 | Stmn1 |
| 4 | Plpp1 | Cyp26b1 | Syne1 | Gja5 | Gbp4 | Plac8 | Hsp90aa1 | Hey1 | Fabp4 | Mki67 |
| 5 | Cmtm8 | Macf1 | Hsph1 | Tgfb2 | Bst2 | Crip1 | Zfp36 | Oaz2 | Gpihbp1 | Ube2c |
| 6 | Cavin2 | F8 | Ppp1r10 | Igfbp3 | Iigp1 | Thbd | Hspa1b | Stmn2 | Akap12 | H2ac24 |
| 7 | Ehd3 | Nostrin | Adamts1 | Slc45a4 | B2m | Mal | Fos | Cxcr4 | Odc1 | H1f5 |
| 8 | Srgn | Hecw2 | Hsp90aa1 | Depp1 | Ly6c1 | Ifi27l2a | Junb | Gng11 | Cxcl1 | H2ac8 |
| 9 | Selenom | Kdr | Phlda1 | Fbln5 | Tapbp | Slc6a6 | Dusp1 | Efna1 | Adamts4 | Prc1 |
| 10 | Cd300lg | Pnisr | Hspa1b | Tm4sf1 | Col4a1 | Npnt | Gadd45b | Ivns1abp | Trp53i11 | Top2a |
| 11 | H2az1 | Psd3 | Btg2 | Ccn2 | Ifitm3 | Tm4sf1 | Hsph1 | Fam174b | Cd74 | Cks2 |
| 12 | Gmfg | Ehd3 | Sox4 | Tmsb4x | Sema7a | Vim | Adamts1 | Hilpda | Rhoc | H1f4 |
| 13 | Cst3 | Ankrd44 | Jun | Cd93 | H2-Ab1 | Adgrl4 | Btg2 | Dll4 | AA467197 | Cenpe |
| 14 | Tagln2 | Arglu1 | Fosb | Esm1 | Gm4951 | Irx3 | Socs3 | Lifr | Inhbb | Cdk1 |
| 15 | Bvht | Ttc14 | Sbno2 | Gja4 | Wars | Glul | Hspa1a | Atp13a3 | Actg1 | Ckap2 |
| 16 | Ddit4l | Tshz2 | Slc38a2 | Sema3g | Col4a2 | Plvap | Phlda1 | Vegfa | Rsad2 | Pclaf |
| 17 | Ramp2 | Sema5a1 | Clk1 | Ccn1 | Serinc3 | Thbs2 | Sertad1 | Dkk2 | Pdgfa | Smc4 |
| 18 | S100a1 | Ddx17 | Apold1 | Jag1 | H2-T23 | Serpina1b | Fosb | Efnb2 | Jam2 | Birc5 |
| 19 | Ddah2 | Dlc11 | Malat1 | Igfbp4 | Gbp9 | S100a6 | Hspb1 | Cyp4b1 | Gbp2 | Smc2 |
| 20 | Ndufa8 | Atrx | Neat1 | Mecom | H2-Aa | Emp3 | Icam1 | Mospd2 | Upp1 | Ccnb2 |
| 21 | Ppia | Igfbp51 | Psd3 | S100a4 | H2-D1 | Calcrl | Plk2 | Plpp3 | Kit | Nav2 |
| 22 | Gimap4 | Nbea | Hbegf | Egfl8 | Emilin1 | Fxyd6 | Ier3 | Igfbp4 | Tubb6 | Nusap1 |
| 23 | Emcn | Srrm2 | Ier5l | Cldn5 | Gbp7 | Sox17 | Ppp1r15a | Unc5b | Cldn5 | Atad2 |
| 24 | Cox4i1 | Ogt | Fus | Pde3a | Psmb9 | Fbln2 | Klf4 | Ctsl | Pde4b | Tubb5 |
| 25 | Crip2 | Shank3 | Son | Stmn2 | Psmb8 | Ehd4 | Nfkbiz | Calm2 | Rnd1 | Kif20b |
| 26 | Nrp1 | Sapcd11 | Hspa1a | Alox12 | Psme1 | Tinagl1 | Jun | Sox18 | Mcam | Cenpa |
| 27 | Cd9 | Arap2 | Gm47283 | Gadd45g | Irf7 | Mest | Apold1 | Pdia5 | Prss23 | Diaph3 |
| 28 | Sh3bgrl3 | Peg3 | Iigp1 | Ntn4 | F8 | Ahnak | Ubc | Apoe | Pkm | Ckap2l |
| 29 | Ppdpf | Ushbp1 | Ppp1r15a | St8sia6 | Col15a1 | Ly6a | Cebpd | Notch1 | Mt1 | Lockd |
| 30 | Dynll1 | Prpf4b | Jund | Ltbp4 | Ly6e | Clic5 | Dnaja1 | Maged2 | Col4a1 | H3c3 |
| 31 | S100a13 | Hipk2 | Macf1 | Efna1 | H2-K1 | Cd93 | Cdkn1a | Tmem176a | Cdkn1a | Pimreg |
| 32 | Pi16 | Rbms1 | Paxbp1 | Vegfc | Slc43a3 | Rdx | Jund | Gstm1 | Actb | Kif23 |
| 33 | Icam2 | Arhgap45 | Atrx | Gad1 | Rtp4 | Edn1 | Hspa8 | Tmem176b | Litaf | H2az1 |
| 34 | Gngt2 | Luc7l3 | Nufip2 | Avpi1 | Psmb10 | Rps3a1 | Ier2 | Gja4 | Nfkbia | Hmmr |
| 35 | Ltc4s | Myo18a | Tra2a | Cavin3 | Stat1 | Pltp | Hbegf | Rgs3 | Tnfrsf12a | Knl1 |
| 36 | Aqp1 | Nsd1 | Lars2 | Atf3 | Xaf1 | Cttnbp2 | Ier5 | Atp1b3 | Sema7a | Tpx2 |
| 37 | Plscr1 | Meis2 | Nbeal1 | Nuak1 | Isg15 | Gas6 | Irf1 | Bnip3l | Smad1 | H2ac15 |

| | | | | | | | | | | |
|---|---|---|---|---|---|---|---|---|---|---|
| **38** | Atp5h | Adgre5 | Klf4 | Prdx4 | Lgals3bp | Rpl3 | Bag3 | Hes1 | Ifrd1 | H1f10 |
| **39** | Atox1 | Numb | Srsf5 | Fbxl7 | Slfn2 | Timp3 | Klf2 | Id2 | Cd93 | Lmnb1 |
| **40** | Tmem176a | Syne2 | Prpf4b | Pcsk5 | Ifi202b | Tmsb4x | Marcksl1 | Gimap5 | Marcksl1 | Aspm |
| **41** | Pde6d | Atp8a1 | Mycbp2 | Klf2 | Tinagl1 | Irx5 | Rhob | Marcks | Col4a2 | Selenoh |
| **42** | Tcn2 | Mapt | Coq10b | Fbln2 | Igtp | Rflnb | Rnd3 | Cdkn1c | Tmsb4x | Hmgn2 |
| **43** | Fam167b | Hmbox1 | Stard9 | Dkk2 | Ifi47 | Rpl9 | H3f3b | Mgll | Rps2 | H1f3 |
| **44** | Ramp3 | Pnn | Meis2 | Bmp4 | Xdh | Rpl8 | Sbno2 | Id3 | N4bp3 | Hjurp |
| **45** | Hsp90b1 | Arhgef15 | Tra2b | Sox17 | Sema3f | Dnm3 | Chka | Mecom | Cd63 | Tmpo |
| **46** | Sumo2 | Ankrd12 | Egr1 | Tsc22d1 | Ifi44 | Sorbs2 | Neurl3 | Gimap1 | Gja1 | Dek |
| **47** | Cox6c | Raph1 | Zfand5 | Ly6e | Il18bp | Ebf1 | Btg1 | Eva1b | Lrrc32 | Anln |
| **48** | Ndufb11 | Nisch | Adamts5 | Actg1 | Irgm1 | Rps5 | Hexim1 | Cbfa2t3 | Bst2 | Incenp |
| **49** | Sod1 | Hdac7 | Mat2a | Slc3a2 | Col18a1 | Rps7 | Hes1 | Tbxa2r | Cebpb | Spc24 |
| **50** | Klk8 | Etl4 | Tle4 | Col18a1 | Lrg1 | Eef1a1 | Tsc22d1 | Rnf7 | Vim | Trim59 |

**Supplementary Table S3: Gene Set Enrichment Analysis of the regenerative EC cluster**

Gene set enrichment analyses of the *regenerative EC* cluster depicting Gene Ontology (GO) terms and corresponding ID of GO:BP MSigDB mouse collections.

| Rank | **regenerative EC** | |
|---|---|---|
| 1 | GO:0048646 | anatomical structure formation involved in morphogenesis |
| 2 | GO:0043408 | regulation of MAPK cascade |
| 3 | GO:0001525 | angiogenesis |
| 4 | GO:0000165 | MAPK cascade |
| 5 | GO:0008219 | cell death |
| 6 | GO:0070371 | ERK1 and ERK2 cascade |
| 7 | GO:0033554 | cellular response to stress |
| 8 | GO:0051094 | positive regulation of developmental process |
| 9 | GO:0006950 | response to stress |
| 10 | GO:0019538 | protein metabolic process |
| 11 | GO:0008283 | cell population proliferation |
| 12 | GO:0009888 | tissue development |
| 13 | GO:0010941 | regulation of cell death |
| 14 | GO:0042981 | regulation of apoptotic process |
| 15 | GO:0043067 | regulation of programmed cell death |
| 16 | GO:0048518 | positive regulation of biological process |
| 17 | GO:0048522 | positive regulation of cellular process |
| 18 | GO:0045765 | regulation of angiogenesis |
| 19 | GO:1901342 | regulation of vasculature development |
| 20 | GO:0006986 | response to unfolded protein |
| 21 | GO:0035966 | response to topologically incorrect protein |
| 22 | GO:0051128 | regulation of cellular component organization |
| 23 | GO:0048514 | blood vessel morphogenesis |
| 24 | GO:0006457 | protein folding |
| 25 | GO:0060548 | negative regulation of cell death |
| 26 | GO:0061077 | chaperone-mediated protein folding |
| 27 | GO:0001568 | blood vessel development |
| 28 | GO:0032101 | regulation of response to external stimulus |
| 29 | GO:0035239 | tube morphogenesis |
| 30 | GO:0006915 | apoptotic process |
| 31 | GO:0012501 | programmed cell death |
| 32 | GO:0050793 | regulation of developmental process |
| 33 | GO:0033043 | regulation of organelle organization |
| 34 | GO:0001944 | vasculature development |
| 35 | GO:0045597 | positive regulation of cell differentiation |
| 36 | GO:0043066 | negative regulation of apoptotic process |
| 37 | GO:0043069 | negative regulation of programmed cell death |
| 38 | GO:0007159 | leukocyte cell-cell adhesion |

| | | |
|---|---|---|
| 39 | GO:1903037 | regulation of leukocyte cell-cell adhesion |
| 40 | GO:0048856 | anatomical structure development |
| 41 | GO:0010942 | positive regulation of cell death |
| 42 | GO:1901564 | organonitrogen compound metabolic process |
| 43 | GO:0009896 | positive regulation of catabolic process |
| 44 | GO:0035295 | tube development |
| 45 | GO:0018130 | heterocycle biosynthetic process |
| 46 | GO:0019438 | aromatic compound biosynthetic process |
| 47 | GO:0034654 | nucleobase-containing compound biosynthetic process |
| 48 | GO:1901362 | organic cyclic compound biosynthetic process |
| 49 | GO:0010563 | negative regulation of phosphorus metabolic process |
| 50 | GO:0042326 | negative regulation of phosphorylation |
| 51 | GO:0045936 | negative regulation of phosphate metabolic process |
| 52 | GO:0045944 | positive regulation of transcription by RNA polymerase II |
| 53 | GO:0009605 | response to external stimulus |
| 54 | GO:0072359 | circulatory system development |
| 55 | GO:0032502 | developmental process |
| 56 | GO:0042127 | regulation of cell population proliferation |
| 57 | GO:0051173 | positive regulation of nitrogen compound metabolic process |
| 58 | GO:0044260 | cellular macromolecule metabolic process |
| 59 | GO:0080134 | regulation of response to stress |
| 60 | GO:0007275 | multicellular organism development |
| 61 | GO:0031328 | positive regulation of cellular biosynthetic process |
| 62 | GO:0006357 | regulation of transcription by RNA polymerase II |
| 63 | GO:0048731 | system development |
| 64 | GO:0031325 | positive regulation of cellular metabolic process |
| 65 | GO:0045893 | positive regulation of transcription, DNA-templated |
| 66 | GO:1902680 | positive regulation of RNA biosynthetic process |
| 67 | GO:1903508 | positive regulation of nucleic acid-templated transcription |
| 68 | GO:0051246 | regulation of protein metabolic process |
| 69 | GO:0043068 | positive regulation of programmed cell death |
| 70 | GO:0032501 | multicellular organismal process |
| 71 | GO:0051716 | cellular response to stimulus |
| 72 | GO:0009891 | positive regulation of biosynthetic process |
| 73 | GO:0006928 | movement of cell or subcellular component |
| 74 | GO:0006366 | transcription by RNA polymerase II |
| 75 | GO:0010557 | positive regulation of macromolecule biosynthetic process |
| 76 | GO:0051247 | positive regulation of protein metabolic process |
| 77 | GO:0071704 | organic substance metabolic process |
| 78 | GO:2000112 | regulation of cellular macromolecule biosynthetic process |
| 79 | GO:0043170 | macromolecule metabolic process |
| 80 | GO:0006807 | nitrogen compound metabolic process |
| 81 | GO:0019220 | regulation of phosphate metabolic process |
| 82 | GO:0042325 | regulation of phosphorylation |
| 83 | GO:0051174 | regulation of phosphorus metabolic process |
| 84 | GO:0002376 | immune system process |

| | | |
|---|---|---|
| 85 | GO:0010556 | regulation of macromolecule biosynthetic process |
| 86 | GO:0006355 | regulation of transcription, DNA-templated |
| 87 | GO:1903506 | regulation of nucleic acid-templated transcription |
| 88 | GO:2001141 | regulation of RNA biosynthetic process |
| 89 | GO:0034645 | cellular macromolecule biosynthetic process |
| 90 | GO:0048870 | cell motility |
| 91 | GO:0051674 | localization of cell |
| 92 | GO:0044238 | primary metabolic process |
| 93 | GO:0044249 | cellular biosynthetic process |
| 94 | GO:0009059 | macromolecule biosynthetic process |
| 95 | GO:0050794 | regulation of cellular process |
| 96 | GO:0031326 | regulation of cellular biosynthetic process |
| 97 | GO:0035556 | intracellular signal transduction |
| 98 | GO:0009653 | anatomical structure morphogenesis |
| 99 | GO:1901576 | organic substance biosynthetic process |
| 100 | GO:0042221 | response to chemical |
| 101 | GO:0006351 | transcription, DNA-templated |
| 102 | GO:0032774 | RNA biosynthetic process |
| 103 | GO:0097659 | nucleic acid-templated transcription |
| 104 | GO:0009893 | positive regulation of metabolic process |
| 105 | GO:0032268 | regulation of cellular protein metabolic process |
| 106 | GO:0010604 | positive regulation of macromolecule metabolic process |
| 107 | GO:0051254 | positive regulation of RNA metabolic process |
| 108 | GO:0051171 | regulation of nitrogen compound metabolic process |
| 109 | GO:0048869 | cellular developmental process |
| 110 | GO:0050896 | response to stimulus |
| 111 | GO:0044271 | cellular nitrogen compound biosynthetic process |
| 112 | GO:0080090 | regulation of primary metabolic process |
| 113 | GO:0009058 | biosynthetic process |

### Supplementary Table S4: Marker genes for spatial transcriptomics

Top differentially expressed genes of single-cell RNA sequencing data for all but the *migrating EC* cluster, used to filter spatial transcriptomics data for cells with similar transcriptional pattern.

| healthy EC | remodeling EC | afferent arteriole EC | injured EC | efferent arteriole EC | regenerative EC | JGA EC | transition EC | proliferating EC |
|---|---|---|---|---|---|---|---|---|
| Ehd3 | Plk2 | Sox17 | Bst2 | Calca | Egr1 | Gng11 | Pdgfa | Cenpf |
| Kdr | Jun | Gja5 | Col4a1 | S100a4 | Dnajb1 | Efna1 | Odc1 | Ube2c |
| | Btg2 | Kdr | Ly6a | Calcrl | Plk2 | Plat | Pde4b | Kdr |
| | Kdr | | Serpina3g | | Kdr | | Col4a1 | |
| | | | Kdr | | | | Kdr | |

**Supplementary Table S5: Clinical renal function parameters in patients:** Summarizes clinical renal function parameters in patients with minimal change glomerular nephritis, complement-mediated TMA, and malignant hypertension-associated TMA; values are presented as mean (M) and standard deviation (SD), and asterisks indicate statistical significance at ✱=p < 0.01.

| | | MCGN | TMA, complement mediated | TMA, malignant hypertension |
|---|---|---|---|---|
| **Gender [n]** | male | 4 | 3 | 6 |
| | female | 4 | 5 | 3 |
| **Age** | M | 40.9 | 44.8 | 45.9 |
| | SD | 20.26 | 15.95 | 12.49 |
| **Creatinin [mg/dl]** | M | 1.07 | 6.91* | 3.54* |
| | SD | 0.81 | 2.76 | 1.51 |
| **Proteinuria [g/d]** | M | 8.56 | 2.83 | 2.26 |
| | SD | 5.43 | 1.84* | 1.48* |
| **Diabetes [0=no/1=yes]** | Yes/No | 0/8 | 0/8 | 0/9 |
| **Hypertonus [0=no; 1=yes]** | Yes/No | 1/5 | 4/0 | 9/0 |
| **Number of glomeruli/section [n]** | M | 22.38 | 28.5 | 27.44 |
| | SD | 9.68 | 24.2 | 8.96 |
| **Global sclerosis [%]** | M | 5.53 | 11.65 | 15.3* |
| | SD | 6.33 | 14.62 | 9.7 |
| **Segmental sclerosis [%]** | M | 0 | 4 | 3 |
| | SD | 0 | 9.03 | 5.77 |
| **Interstitial Fibrosis and Tubular Atrophy [%]** | M | 4 | 15 | 31.1* |
| | SD | 1.65 | 15.49 | 15.42 |
| **Inflammation [0=no; 0,5=minimal; 1=mild; 1.5=mild-moderate; 2=moderate; 3=severe]** | M | 0 | 1* | 0.9* |
| | SD | 0.35 | 0 | 0.37 |
