## Supplementary Methods for "Spatiotemporal Mapping of Glomerular Endothelial Regeneration Identifies *Apold1* as an Intrinsic Regulator of Repair"

**Organ Collection and Tissue Processing**

On the day of sacrifice, mice were anesthetized with inhaled isoflurane (2%) and transcardially perfused with 0.9% (w/v) NaCl to remove blood components from both kidneys. Transcardial perfusion with saline/PBS is commonly used to clear blood from tissues prior to organ collection or fixation The left kidney was harvested, bisected, and fixed overnight at 4°C in zinc-based fixative, methanol-Carnoy fixative, or 4% paraformaldehyde (PFA). Fixed kidney tissues were subsequently processed and embedded in paraffin for further histological analyses.

**Immunohistochemistry using human kidney biopsies**

The use of kidney biopsy material has been approved by the Ethics Committee of the Friedrich-Alexander University of Erlangen-Nuremberg, waiving the need for retrospective consent for the use of archived excess material (Ref. No. 22-150-D). Formalin-fixed, paraffin-embedded sections from human kidney biopsies were deparaffinized and rehydrated followed by blocking of endogenous peroxidase activity with 3% H2O2 diluted in deionized water and incubated for 10 min at room temperature. Antigen retrieval was performed in target retrieval solution pH 6 (DAKO Deutschland, Hamburg, Germany) for 2.5 min in a pressure cooker (HIER). For IHC staining, an avidin-biotin blocking kit (Vector Laboratories, Burlingame, CA, USA) was applied and sections were layered with the primary antibodies previously validated and published in the Human Protein Atlas (www.proteinatlas.org; Supplementary table S5). After removal of unbound primary antibody by washing with wash buffer (50mM Tris pH 7.4 supplemented with 0.1% (v/v) Tween 20), biotinylated secondary antibodies (biotinylated anti-mouse IgG or biotinylated anti-goat IgG, Vector Laboratories, 1:500) was added and incubated for 30 min at room temperature (RT). The ABC-Kit and Vectastain DAB Immpact Kit (both Vector Laboratories) were used to visualize the bound biotinylated secondary antibody. Each slide was counterstained with hematoxylin. For negative controls, immunostaining was performed by omission of the primary antibody.

**Xenium Spatial Transcriptomic Analysis**

For the spatial transcriptomic analysis shown in Figure 5, cell segmentation was performed using the 10x Genomics Xenium workflow. Briefly, individual transcript molecules were assigned to spatial coordinates within the tissue section, followed by morphology-based cell segmentation to define cell boundaries and assign transcripts to individual cells.

Glomerular regions were identified based on renal tissue morphology and spatial localization. Within these regions, glomerular endothelial cells were annotated according to the expression of endothelial and glomerular endothelial marker genes, including Ehd3 and Plat. Segmented cells located within glomerular regions and expressing these markers were included in downstream analyses of glomerular endothelial cell-associated gene expression.
